# Identification of sex-determining loci in hybridizing *Catostomus* fish species

**DOI:** 10.1101/2022.05.03.490509

**Authors:** Cassandre B. Pyne, S. Eryn McFarlane, Elizabeth G. Mandeville

**Affiliations:** Department of Integrative Biology, University of Guelph, Guelph, Ontario, Canada; Department of Botany, University of Wyoming, Laramie, Wyoming, United States

**Keywords:** Sex determination, sex chromosomes, *Catostomus*, genotyping-by-sequencing, GEMMA

## Abstract

Despite the near-universality of gonochorism (separate sexes) in eukaryotic organisms, the underlying mechanisms of sex determination are poorly understood and highly variable in some taxa. Sex determination mechanisms may promote or impede reproductive isolation depending on whether mechanisms are similar between related species, so identifying genomic regions associated with sex determination is especially relevant for understanding evolutionary diversification and hybridization between closely related species. In *Catostomus* fishes, contemporary hybridization is variable and extensive, but factors influencing hybridization dynamics are not fully understood. In the present study, we aim to describe the genetic basis of sex determination in bluehead (*Catostomus discobolus*) and white suckers (*Catostomus commersonii* ) to understand the potential impact of sex determination on reproductive isolation. We used genotyping-by-sequencing data from *Catostomus* species and their hybrids to identify regions of the genome associated with sex using a genome-wide association study and the identification of sex-specific loci. We identified a genetic basis of sex determination in *Catostomus* fishes, with a region of the genome significantly associating with sex in bluehead suckers. This region is suggestive of a master sex-determining region in bluehead suckers but is not significant in white suckers, implying that either the sex-determining region of the genome differs in these two species that hybridize, or that sample size was insufficient to identify this genomic region in white suckers. By describing and comparing sex-determination systems across *Catostomus* fish species, we highlight the relationship between sex determining systems and hybridization in closely related fish species.

## 2 Introduction

Sex determination is the system that determines whether an individual will be male or female and is surprisingly diverse in many clades of plants and animals (Ashman *et al*. 2014). Regions of the genome associated with sex determination are hypothesized to have played a major role in evolutionary diversification and speciation (Payseur *et al*. 2018). Incompatible sex-determination systems can be an effective reproductive barrier that can restrict gene flow and may play a role in hybrid sterility (Haldane 1922, Hubbs 1955, Delph & Demuth 2016, O’Neill & O’Neill 2018). Not only the systems themselves, but the rapid evolutionary turnover of sex determination in some species, can impact speciation (Kitano & Peichel 2012). However, we cannot understand the role sex determination plays in regulating reproductive isolation and evolutionary diversification without understanding the mechanisms that underlie it. Remarkably, in many non-model organisms, genetic mechanisms of sex determination remain completely unknown.

Genetic mechanisms of sex determination in gonochoristic species are variable and can range from large heteromorphic sex chromosomes, where sex chromosomes are morphologically differentiated, to multiple sex-associated loci (Mank *et al*. 2006), represented by unlinked genes across the genome or localized regions with many sex-determining genes (Kobayashi *et al*. 2013, Heule *et al*. 2014). Master sex-determining genes control sexual differentiation by acting as a switch that leads to either the male or female phenotype (Yano *et al*. 2012, Heule *et al*. 2014). However, as more studies use genome-wide data to interrogate sex determination systems, there is increasing evidence for polygenic sex determination, where sex is controlled by multiple, possibly interacting loci (Vaneputte *et al*. 2007). Recent work suggests that sex determination is more complex and polygenic than originally thought (Bachtrog *et al*. 2014, Heule *et al*. 2014, Roberts *et al*. 2016, Faggion *et al*. 2019, Adhikari *et al*. 2021).

Sex-determination systems are particularly variable in fish. The genetic elements that control sex determination in fish are less conserved than in some other vertebrate clades like mammals and birds, suggesting multiple evolutionary transitions between sex-determining systems in fish (Heule *et al*. 2014). Both genetic and environmental sex determination have been documented in fish; however, evidence is accumulating that sex determination systems that rely on multiple genetic elements, potentially with environmental influence, are more frequent than initially recorded (Heule *et al*. 2014). The genetic architecture of sex determination in fish can vary beyond heteromorphic sex chromosomes. For example, sex can be determined with just one locus or gene such as *SRY* or *dmrt1b*, or multiple loci such as *sar3*, *sar5* and *sar16* in zebrafish (Anderson *et al*. 2012). In sea bass, both genetic factors and temperature influence the sex of the individual (Palaiokostas *et al*. 2015). This is also seen in Nile tilapia (*Oreochromis niloticus*), channel catfish (*Ictaluras punctatus)*, and Atlantic salmon (*Salmo salar* ) (Baroiller *et al*. 2009, Eisbrenner *et al*. 2014, Kijas *et al*. 2018, Bao *et al*. 2019). Many fish species do not have a sex determination system that relies on a known master sex-determining gene which makes the mechanisms of sex determination even more difficult to detect (Palaiokostas *et al*. 2015, Faggion *et al*. 2019). This leaves a large gap in knowledge on the underlying genetics and mechanisms of sex determination, particularly for non-model fish taxa. Not only are the mechanistic pathways and specific genes involved in sex determination mostly unknown, but overall systems are undescribed in many teleost fish species (Mank *et al*. 2006).

Species with historical and contemporary hybridization, such as *Catostomus* fishes, can shed light on the role of sex determination in reproductive isolation and hybridization outcomes, as mismatching sex determining systems can lead to strong post-zygotic isolation (Haldane 1922, Coyne & Orr 2004, Kitano & Peichel 2012, Lima 2014). *Catostomus* fishes (“suckers”) are a genus of freshwater fishes that are widespread through North America. *Catostomus* fishes are historical tetraploids, as they diversified following an allopolyploid genome duplication event approximately 50 million years ago at the base of the Catostomidae family (Hubbs 1955, Uyeno & Smith 1972). Hybridization events followed diversification from the allopolyploid ancestor, leading to the species richness and diversity seen in the *Catostomus* genus. In total, there are 26 species including both native and introduced species (Dowling *et al*. 2016). Most species in the Catostomidae family have approximately 100 chromosomes, with an absence of well-differentiated sex chromosomes (Uyeno & Smith 1972). Some species of *Catostomus* suckers are of conservation concern, such as the bluehead and flannelmouth sucker (Cooke *et al*. 2005). Additionally, suckers do not display obvious sexual dimorphism; thus, identifying males and females can only be achieved during the spawning season.

*Catostomus* fishes experience extensive contemporary hybridization between both native and introduced species. Hybridization is incredibly variable and can be human induced through the introduction of species such as the white sucker (Mandeville *et al*. 2015, 2017). Hybridization in *Catostomus* fishes in the Upper Colorado River basin primarily involves three species, bluehead (*Catostomus discobolus*), flannelmouth (*C. latipinnis*), and white sucker (*C. commersoni* ). Hybridization outcomes vary from F1 hybrids to extensive backcrossing depending on the location or species pair. For example, hybridization between the flannelmouth and white sucker can produce later-generation hybridization including backcrossing to both parental species, while bluehead×white hybrids are mostly F1 hybrids (Mandeville *et al*. 2017). Additionally, flannelmouth×white hybrids have asymmetrical backcrossing depending on the location, where some locations have hybrid backcrossing to only one parental species (Mandeville *et al*. 2017). The variation in hybridization and reproductive isolation makes *Catostomus* fishes a unique study system to investigate the genetic basis of sex determination.

There is a possibility that sex determination incompatibilities may play a role in the variation in reproductive isolation across hybrid crosses in *Catostomus* fishes. For example, the hybrid that results from reproduction between bluehead and white suckers has limited backcrossing to the parental species which indicates some level of reproductive isolation between these species, although the mechanism is unknown (Mandeville *et al*. 2015). Therefore, sex determination may be an effective mechanism driving reproductive isolation between the hybrid and parental species. Sex determination is mostly undescribed in *Catostomus* suckers however, other fish in the Cypriniformes order show some form of genetic sex determination including male and female heterogamety (Ashman *et al*. 2014). Therefore, there may be a genetic basis in suckers (Bachtrog *et al*. 2014). The potential for sex determination to be a driver for reproductive isolation in *Catostomus* suckers is an intriguing possibility and makes these fish an appropriate model in which to investigate the role of sex determination.

In this study, we investigated the genetic basis of sex determination in bluehead suckers (*C. discobolus*), in addition to a smaller sample of white suckers (*C. commersonii* ) and bluehead×white hybrids. We used genomic data from *Catostomus* species and their hybrids to determine regions of the genome that were significantly associated with sex. We investigated both sex-associated loci (those present in both sexes but differentiated) and sex-specific loci (those that are exclusively in one sex). Sex-associated loci are consistent with a sex-determining region, polygenic sex determination, or homomorphic sex chromosomes, where sex chromosomes are morphologically indistinguishable. Sex-specific loci correspond to heteromorphic sex chromosomes. A secondary aim of this study was to determine to what extent the genetic basis of sex determination is variable across *Catostomus* sucker species. Due to the high evolutionary turnover of sex-determining mechanisms in fish, all species of *Catostomus* suckers may not share the exact same underlying genetic mechanisms. If mechanisms differ across species, it may explain some of the variability in hybridization outcomes across species pairs in *Catostomus* fishes.

## 3 Materials and methods

### Sampling

We used existing data from fish collected for a previous sequencing effort (n=139, Mandeville et al., unpublished data) as well as new data collected for this study (n=130). The data are from *Catostomus* sucker fin clip samples from sampling in 2015 and 2019 in the Gunnison River basin (Colorado, USA), Jackson Lake (Wyoming, USA), and small streams in southern Ontario (Canada). The total data set consists of 269 individuals of known sex and includes three species, *C. discobolus* (bluehead sucker, n=155), *C. latipinnis* (flannelmouth sucker, n=12), and *C. commersonii* (white sucker, n=46) as well as their hybrids (Table 1, Table S1, Table S2). Each individual’s sex was determined by experienced field personnel by visual inspection during spawning season or by dissection.

**Table 1:** Number of individuals per species or hybrid and their sex. Males, females, and probable females (those that could not be conclusively sexed as female) are included.

| **Species** | **Males** | **Females** | **Probable Females** | **Total** |
| --- | --- | --- | --- | --- |
| Full data set | 131 | 87 | 51 | 269 |
| Bluehead sucker | 73 | 37 | 45 | 155 |
| White sucker | 33 | 9 | 4 | 46 |
| Bluehead×white hybrid | 15 | 33 | 0 | 48 |

### Sample preparation and sequencing

We completed DNA extractions on the fin clips using the DNeasy Blood and Tissue Kit according to the manufacturer’s instructions (Qiagen, Inc) and quantified the samples with a Nanodrop spectrophotometer. The extracted DNA was used to create a genomic library for DNA sequencing. Specifically, we used reduced-representation genotyping-by-sequencing (GBS) methods, where we used restriction enzymes ‘EcoRI’ and ‘MseI’ to digest genomic DNA into fragments (Parchman *et al*. 2011). Barcodes and identifiying adaptor sequences were ligated to each DNA fragment for amplification. After library preparation, we used a PippinPrep (D-Mark Biosciences) to size select DNA fragments with lengths between 250 and 350 base pairs. The two prepared libraries were then sent to University of Texas Genome Sequencing and Analysis Facility for high-throughput DNA sequencing on an Illumina HiSeq 2500 and Toronto’s The Centre for Applied Genomics at The Hospital for Sick Children for sequencing on an Illumina NovaSeq 6000.

A new flannelmouth sucker (*C. latipinnis*) reference genome was used, which improved upon an artificial reference genome used in a previous study (Mandeville *et al*. 2017). The individual sampled was from Little Sandy Creek, near Farson, Wyoming. This individual was male and was sampled by Wyoming Game and Fish. The high-quality reference genome was produced at Dovetail Genomics using a draft assembly in addition to data from Hi-C and Chicago libraries. The reference genome has an N50 of 37.5 Mb and is assembled into nearly chromosome-level scaffolds (Mandeville et al., unpublished data).

### Variant Calling and Filtering

After sequencing, the resulting .fastq files were parsed to separate out individual fish with differing identifying barcode sequences using a custom `perl` script. Reads for each individual fish were then aligned to the flannelmouth sucker reference genome (Mandeville et al., unpublished data) using `bwa` (version 0.7.17) (Li & Durbin 2009). This resulted in binary alignment map (BAM) files that were then used for variant calling using `samtools` (version 1.12) and `bcftools` (version 1.11) (Li *et al*. 2009, Li 2011). Computing was completed on the Graham cluster associated with Compute Canada, through a Resources for Research Groups (RRG) allocations.

We used `vcftools` (Danecek *et al*. 2011) to filter variants, where we filtered to retain loci that were present in more than 50% of individuals, had a minimum quality of 20 and a minor allele count of at least 3. We also removed individuals with more than 90% missing data. Later stages of filtering screened for a minor allele frequency of 0.05 and restricted single nucleotide polymorphisms (SNP) to one SNP per contig. Restricting contigs to have only one SNP reduces the chance that SNPs are in linkage disequilibrium. In addition, potential paralogs were detected using a custom R script and removed using `vcftools` (Danecek *et al*. 2011).

After filtering, we used `entropy` to determine the ancestry of the hybrids and sort individuals by species (Gompert *et al*. 2014, Shastry *et al*. 2021). `entropy` estimates genetic parameters and individual ancestry using a Bayesian model and Markov Chain Monte Carlo. Sequence data are incorporated into the model using genotype likelihoods (Shastry *et al*. 2021), which were estimated using `bcftools`. To determine the optimal value of k (number of genetic clusters), we ran `entropy` for k=1 to k=6, corresponding to the maximum possible number of *Catostomus* species that could be included in our samples (Mandeville *et al*. 2017). Genotypes and associated likelihoods were used in a principal components analysis (prcomp in R), where k-means clustering and linear discriminant analysis were applied to the results in R (R Core Team 2022) to create starting values of q (in `entropy`, admixture proportion). We used non-random starting values for q because starting in a better parameter space (as informed by the data) speeds model convergence (Shastry *et al*. 2021). We ran three replicates for each value of k. Each replicate included 50,000 MCMC (Markov Chain Monte Carlo) steps, including 25,000 burn-in discarded values, and we kept every 25th value. `entropy`’s output is genotype probability and ancestry estimates which allows for grouping of individuals to genetic clusters.

### Genome-wide association study

We used a genome-wide association study (GWAS) to detect associations between regions of the genome and sex in both sexes. GEMMA, a single-locus software, was used to fit a linear mixed model (LMM), where the effect sizes of SNPs are assumed to be from a normal distribution (Zhou & Stephens 2012). GEMMA only allows loci that have been genotyped in at least 95% of the individuals; therefore, genotypes were imputed using the genotype likelihoods from the output of entropy in a custom R script (Shastry *et al*. 2021). Prior to conducting the association analysis, GEMMA calculates a relatedness matrix to account for population structure. The random effects are expected to be structured by population and species; therefore, including this in the association analysis leads to more reliable results.

To detect loci that are associated with sex and are present in both sexes (i.e., not sex-specific), we ran GEMMA on all 260 individuals (after filtering) of known sex and including all three species and hybrids. Initially, we used GEMMA on the entire data set to investigate whether there was a shared basis of sex determination that emerged among all species and hybrids. We next split the data set by species and hybrid (bluehead, white, bluehead×white) to determine differences in sex determination systems across species. Filtering as outlined above was completed for each species. SNPs were then visualized using Manhattan plots with the `qqman` package in R (version 0.1.8) (Turner 2018). SNP p-values were adjusted using the negative log function, and the significance threshold was corrected using the Bonferroni correction.

In addition to the LMM in GEMMA, we also used the Bayesian sparse linear mixed model (BSLMM). This model uses Markov Chain Monte Carlo to estimate the proportion of phenotypic variance explained by the genotypes (Zhou *et al*. 2013). BSLMMs are beneficial to use in addition to LMMs because they assume that each variant falls under two distributions and does not assume non-zero SNP effects a priori. In the first distribution, all variants are assumed to have a normal distribution of effects, as is the case in LMMs (Zhou *et al*. 2013). In the second (the ‘sparse’ matrix), a small number of variants are assumed to have an effect, as is the case in Bayesian variable selection regression models (BVSR) (Guan & Stephens 2011, Zhou *et al*. 2013). Using both models complement each other because they use different assumptions about how explanatory power is weighted across SNPs. This model was run using the same input as described above. We ran BSLMM with 25 million iterations and a burn in of 10 million iterations. Samples were taken every 100 iterations. White and bluehead×white hybrids, however, did not converge with these parameters; therefore, the model run conditions were adjusted to ensure convergence. White suckers converged with 75 million iterations and a burn in of 15 million, with samples taken every 100 iterations. Bluehead×white hybrids converged with parameters of 1 million iterations with a burn in of 100 thousand, with samples taken every 100 iterations. BSLMM was run three times for each data set to ensure convergence. MCMC distributions of the phenotypic variation (PVE) were used to assess convergence of the model. Results were visualized in R using a GEMMA workshop (Soria-Carrasco 2019). Significance was quantified using posterior inclusion probability (‘PIP’) which indicates the proportion of iterations in which the SNP is included in the sparse matrix, suggesting a non-zero effect size.

### Discriminant analysis of principal components

We also identified candidate sex-associated loci using DAPC (discriminant analysis of principal components, Jombart 2008), implemented through the adegenet package in R (Jombart *et al*. 2010) and following analyses from Junker *et al*. 2020 and Meuser *et al*. 2022. This approach uses a principal components analysis to identify major axes of genetic variation, then uses discriminant analysis with sex as the grouping variable to identify sex-associated loci. We completed analyses separately for bluehead and white suckers. For each data set, we used a randomization procedure to generate a statistical threshold for SNP loading, and identified loci that exceeded the 99.5% quantile as sex-associated for bluehead suckers and 99.9% for white suckers. The threshold for white suckers is higher than bluehead suckers to account for geographic variability in white sucker samples and larger number of SNPs in the data set.

### Additional analyses

To complement the results of the GWAS and DAPC, we also investigated F*_ST_* and linkage disequilibrium for significant loci from the GWAS. Using VCFtools, we calculated the Weir and Cockerham F*_ST_* to assess genetic differentiation between the sexes (Weir & Cockerham 1984, Danecek *et al*. 2011). Significant outliers were detected using Rosner’s test. Additionally, we calculated pairwise *r*^2^ (squared correlation coefficient) to estimate linkage disequilibrium between significant SNPs using VCFtools (Danecek *et al*. 2011).

To determine the effect of sample size on GWAS results, we down-sampled the bluehead (n=150) and entire (n=260) data sets to match the phenotypic classes with fewer individuals. Since the white and bluehead×white sucker datasets have approximately 45 individuals each, we randomly sampled 45 individuals from both the bluehead suckers and entire data sets. We then completed filtering as described above, and imputed any missing genotypes. We then ran both LMM and BSLMM models to investigate any differences between the larger and down-sampled sample size of the bluehead and entire data sets. This process was completed a total of three times each for the bluehead and entire data sets to ensure that the resulting pattern was similar across replicates.

During data collection, 19% of individuals were phenotypically identified as ‘probable female’ rather than either males or females. These individuals could not be conclusively determined to be females, as they were not yet in spawning condition. However, males were in spawning condition; therefore, individuals were marked as ‘probable females’ by expert field personnel familiar with spawning time in this system. To validate that these individuals were female, we used GEMMA’s LMM, BSLMM, and phenotype prediction functionality, which can predict phenotypes using the present genomic data and available phenotypes (see Supplement for more information).

### Sex-specific loci identification

In order to detect regions of the genome that are associated with one sex but not the other, we used the beta version of a custom program `Ms.SSLI` (Pyne and Mandeville, unpublished software available at https://github.com/pynec/Ms.SSLI). This program detects genomic regions associated with heteromorphic sex chromosomes and uses allele depth values to determine loci that are exclusively in one sex using a statistical threshold (Stovall *et al*. 2018). Allele depth is the count of reads that support an allele for an individual and is a standard field in a Variant Call Format file (VCF) (Danecek *et al*. 2011). The input data was derived from the same variant call format file used for GWAS, but we applied different filtering regimens, as the missingness filtering criteria was loosened to allow for the detection of loci that by definition (sex-specific) are expected to have missing data in about half of the individuals. Loci could have up to 90% missing data while individuals were allowed up to 80% missing data. The resulting VCF was then used to extract the allele depth using `vcftools` version 0.1.16 (Danecek *et al*. 2011). RADSex was also used as a complementary technique to identify sex-specific loci (Feron *et al*. 2020). This workflow takes .fastq files of demultiplexed reads and a phenotype file with individuals and sex as input. RADSex will then determine which monomorphic markers are significantly associated with sex through a presence-absence scan and Pearson’s chi-squared test rather than a statistical threshold as in `Ms.SSLI`(Feron *et al*. 2020). Thus, it is able to identify both homomorphic and heteromorphic sex chromosomes (Feron *et al*. 2020). Output was visualized with the `sgtr R` package (Version 1.1.4) (Feron *et al*. 2020).

## 4 Results

DNA sequencing resulted in approximately 1.9 x 10^8^ million raw reads. The average number of raw reads per individual was 1225761 and the mean number of assembled reads was 1115773, leading to an average of 91% of reads aligning to the flannelmouth sucker reference genome (Mandeville et al., unpublished data). From the assembled reads, variants were identified. We removed 9 individuals with more than 90% missing data and removed 4,253 potentially paralogous loci. This resulted in 27,822 SNPs for 260 individuals that were kept after variant calling and filtering. Mean sequence coverage was 5.17 reads per locus per individual. After running entropy, we confirmed that there are three genetic clusters, corresponding to the three species in the data. (Fig. S1). Using the DIC (Deviance Information Criterion) values, the k=3 and k=4 models were best supported and had similar DIC values, but we continued with k=3 as it matches the three phenotypically identified species (Fig. S2). In addition, using a principal components analysis, we confirmed that individuals of the same species cluster together along the first 3 principal components (Fig. S3). The three species and their hybrids formed three distinct genetic clusters across the principal components. PC1 accounts for 98.2% of the variation and separates each of the species.

Following initial analyses with the entire data set, the data were grouped by bluehead suckers, white suckers and bluehead×white hybrids. Filtering resulted in 17,579 loci for 150 bluehead sucker individuals. These data had a mean coverage of 4.55 reads per locus per individual. For white suckers, there were 69,415 loci for 44 individuals with a mean coverage of 5.34 reads per locus per individual. Lastly, filtering of bluehead×white hybrids resulted in 59,295 loci for 46 individuals with a mean coverage of 5.04 reads per locus per individual. Filtering with a relaxed missingness threshold for the identification of sex-specific loci resulted in 49,224 loci with a mean coverage of 3.39 reads per locus per individual for bluehead suckers and 95,864 loci with a mean coverage of 4.45 for white suckers. Bluehead×white hybrids had 85,516 loci with a mean coverage of 4.21. Including ‘probable females’ as females lead to qualitatively similar results as excluding ‘probable females’ suggesting that these individuals are most likely female, as initially identified by expert field personnel (Fig. S4). For the following analyses, probable females were included as females (see Supplement for more information).

### Sex-associated loci in bluehead suckers

We identified sex-associated loci in bluehead suckers. Specifically, we found two sex-associated SNPs that are strongly supported by several analyses (Fig 1). These SNPs were detected using both LMM and BSLMM (LMM: p=10*^−^*^15^, p=10*^−^*^10^; Fig. 2A, BSLMM: PIP=1.0, PIP=0.42, Fig. 2B). Estimates reported from BSLMM are from the first run. Another significant SNP from BSLMM was located on chromosome 19 and had a PIP of 0.35. All three runs of BSLMM converged and consistently demonstrated the same well-supported loci (Table S3). We used the phenotype prediction function to determine whether the loci contributing to sex in bluehead suckers could predict sex in white suckers, and found that these SNPs could not predict sex in white suckers as the predicted phenotype values were variable and 27% of the time did not correspond to the correct sex. Therefore, the identified sex-associated loci most likely have poor predictability in white suckers. No sex-specific loci were detected in bluehead suckers using `Ms.SSLI` or RADSex (Fig. S5). We report p-values and beta estimates for the entire data set, bluehead, and white suckers in Supplementary tables S4, S5, and S6 respectively. We used beta estimates as a proxy for effect size. DAPC results also identify SNPs on chromosome 4 as sex-associated loci in bluehead suckers. Using a SNP loading statistical threshold of 0.995, 55 sex-associated loci were identified across 36 chromosomes in bluehead suckers. 3 sex-associated loci were found on chromosome 4, with one overlapping with the loci identified by LMM and BSLMM (SNP loading=0.000742142, Table S7).

**Figure 1:**
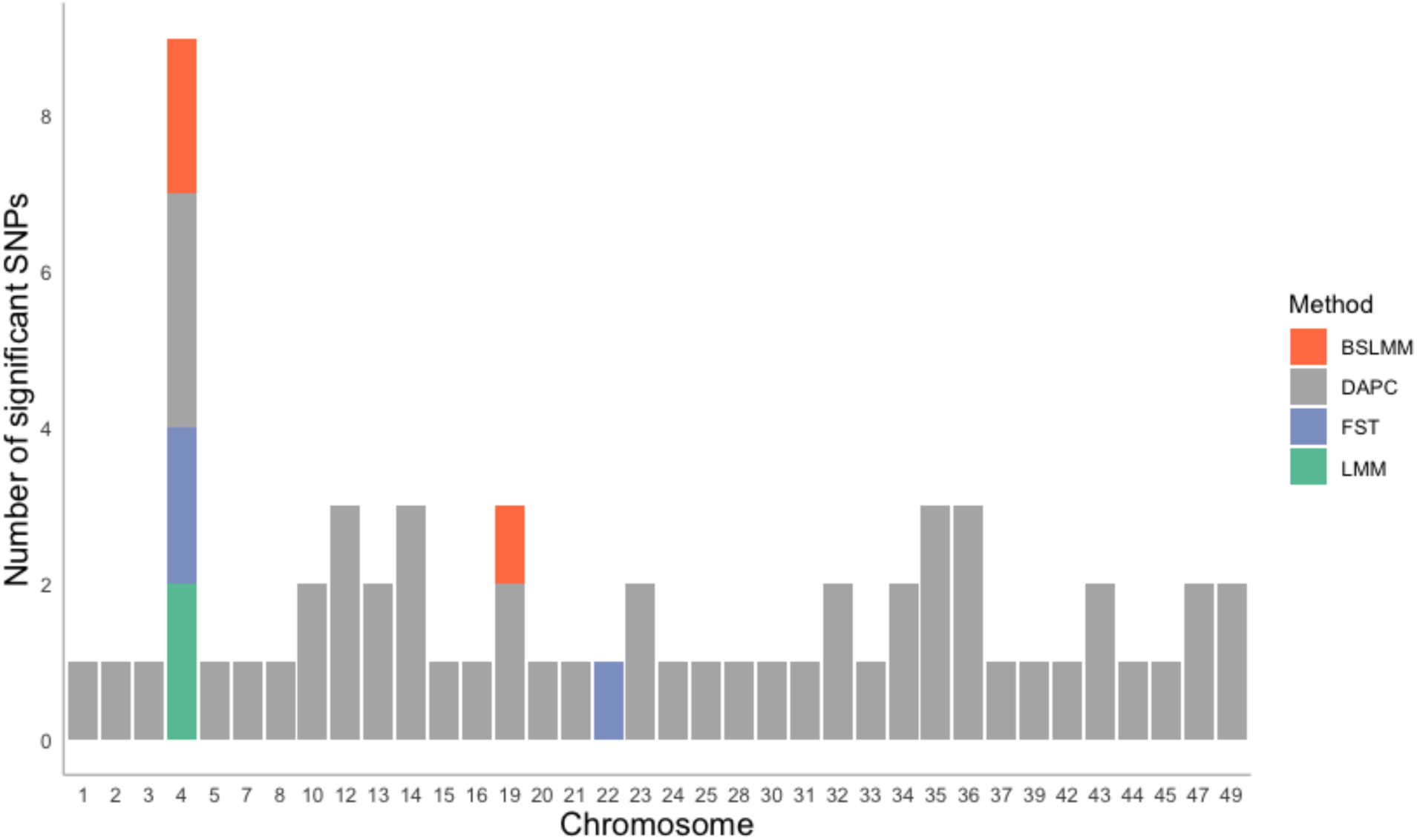
Barplot depicting the overlap of significant SNPs for bluehead suckers (n=150) using GEMMA’s LMM, BSLMM, DAPC, and FST outliers. Only the chromosomes that had significant SNPs are shown on the plot. There is only one SNPs on chromosome 4 (at base pair position 27320056) that is significant in all 4 methods.

**Figure 2:**
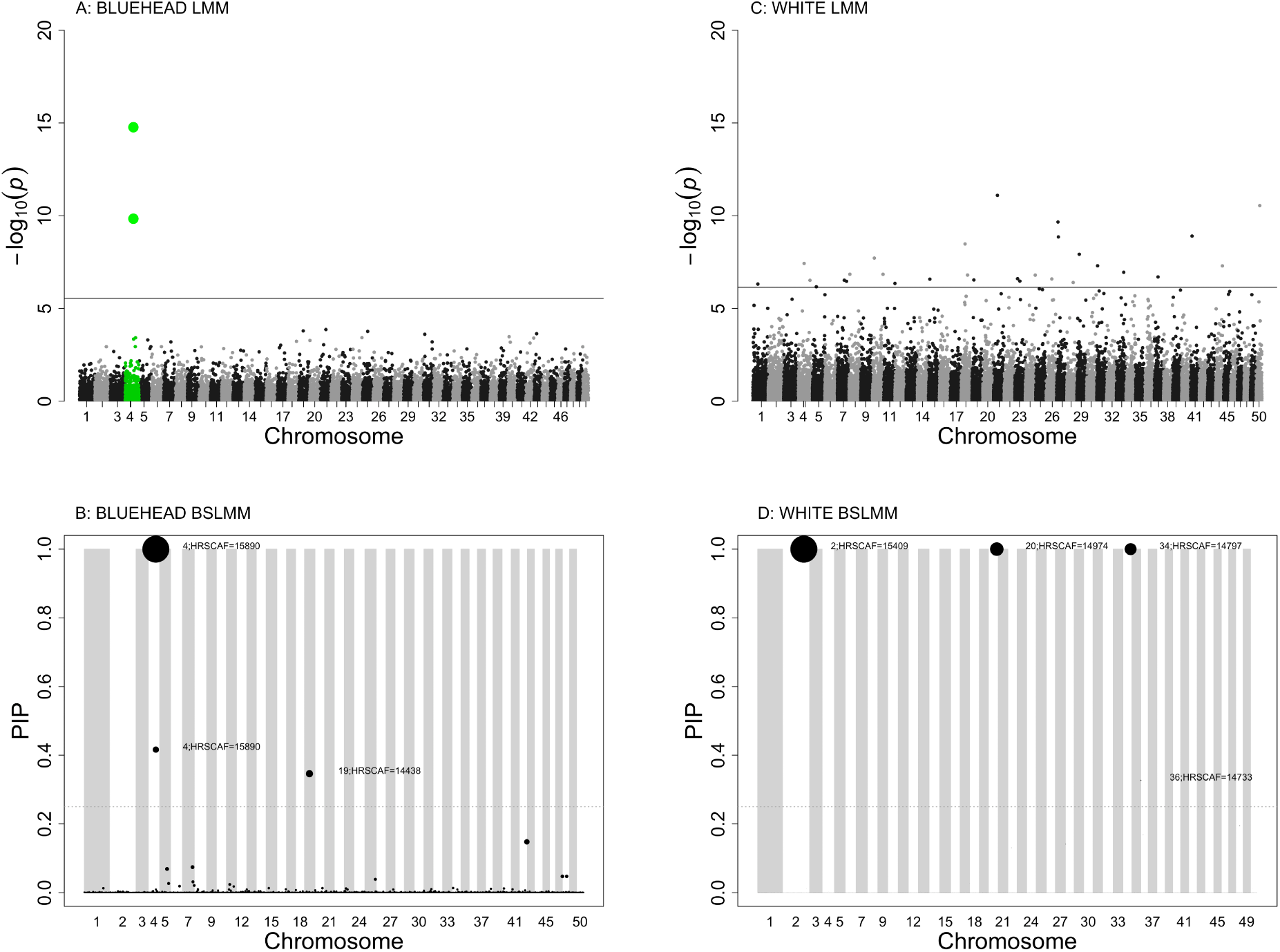
Sex-associated loci in the bluehead and white *Catostomus* fishes datasets across the 50 largest scaffolds relative to the flannelmouth reference genome. a) Manhattan plot from GEMMA’s LMM between SNPs and sex for bluehead fishes (n=150), with a significance threshold of p=2.84*^−^*^6^. The most significant chromosome, chromosome 4, is highlighted in green. b) PIP (posterior inclusion probability) of SNPs across chromosomes from the BSLMM for bluehead suckers (n=150). The dotted line indicates the threshold of PIP value of 0.25, and the grey and white blocks visually show the chromosomes. The sparse effect is indicated by the size of the point. c) Manhattan plot from GEMMA’s LMM between SNPs and sex for white suckers dataset (n=44). The black line indicates the Bonferonni corrected significance threshold of p=1.80*^−^*^6^. d) PIP of SNPs from GEMMA’s BSLMM for the white (n=44) data set. As with b), The dotted line indicates the threshold of PIP value of 0.25 and the sparse effect is indicated by the size of the point.

The two sex-associated loci we identified on chromosome 4 with LMM and BSLMM form a sex-associated haplotype that participates in sex determination in bluehead suckers. The two sex-associated loci are located closely together on chromosome 4, at base pair position 27210425 and 27320056 (Fig. 3). The *r*^2^ value between the two loci is 0.239 where the average along chromosome 4 is 0.048, suggesting that these two loci are linked. Additionally, these two loci are the most divergent between the sexes, with an F*_ST_* of 0.27. One of these loci, at base pair position 27320056 was identified by all four methods (LMM, BSLMM, DAPC, F*_ST_* , Fig. 1). Of the 108 available female genotypes in the sex-associated loci haplotype, 94% of females were homozygous for the same genotype. Of the 82 available male genotypes, 71% were heterozygous for the same genotype. However, when just investigating one of the two sex-associated loci, females were 89% homozygous for the second sex-associated locus while males were 74% heterozygous. We also estimated heritability of these loci using as the proportion of phenotypic variance explained by *V_SNP_* (*V_SNP_* =2*pqa*^2^, where p and q are the major and minor allele frequencies and a is the effect size; (Falconer & Mackay 1996). Heritability for the two significant loci on chromosome 4 was 0.44 and 0.14, respectively.

**Figure 3:**
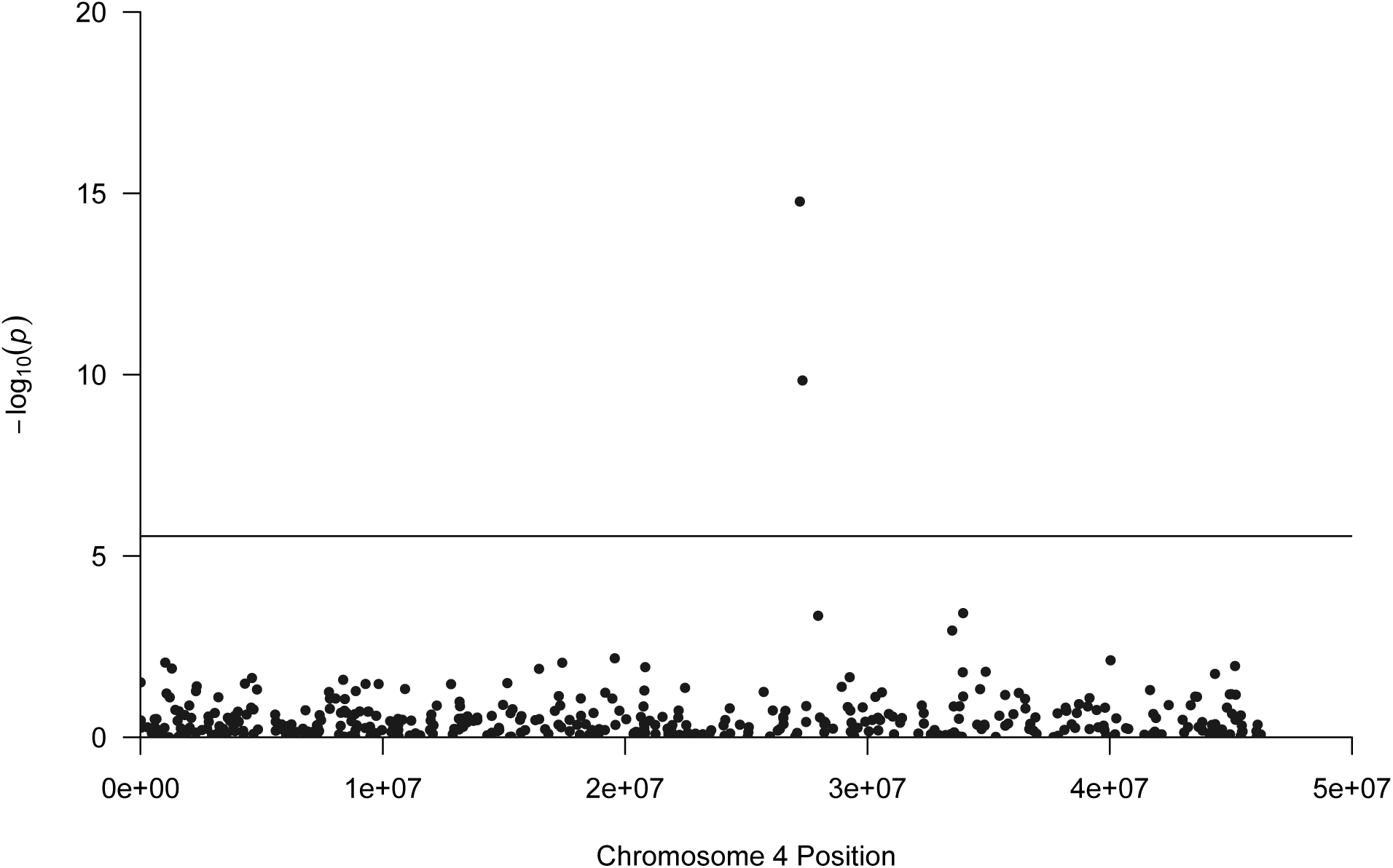
Manhattan plot depicting base pair positions on chromosome 4 and the location of significantly sex-associated SNPs in bluehead suckers (n=150). The significantly sex-associated SNPs shown in the plot are from GEMMA’s LMM. The genome-wide significance threshold shown indicates the Bonferonni corrected significance threshold of p=2.84*^−^*^6^.

We identified 3 sex-associated SNPs across 3 chromosomes when using the entire data set with all species and hybrids (n=260, LMM; Table S5). However, chromosome 4, which had one significant SNP (p=10*^−^*^19^) was the most sex-associated, similar to bluehead suckers (Fig. S6A). This SNP was significant in the LMM and well-supported in the BSLMM (PIP=1.0; Fig. S6B, Table S3). The other two significant SNPs were not well supported using BSLMM (PIP= 0.005, PIP=0.028). The three runs of BSLMM for the entire data set converged and were consistent with reporting the same well-supported loci.

### Evidence for a genetic basis of sex determination in white suckers

Analyses point to a genetic basis of sex determination in white suckers, although the genetic architecture remains unclear, as different loci were significant with different analyses. In contrast to bluehead suckers, there were 29 sex-associated SNPs across 23 of the 50 largest chromosomes in white suckers, indicating a polygenic architecture of sex determination (LMM, Fig. 2C, Table S6). None of these SNPs overlapped with SNPs identified in the full data set or bluehead suckers (Fig. S7). 11 of the 23 chromosomes with significant SNPs had loci with large PIP values in the three BSLMM runs (Fig. 2D, Table S3). However, only chromosome 27 had loci with high PIP values in more than one run of BSLMM. The sex-associated chromosome (not SNP) in blueheads, 4, was also found to be sex-associated in white suckers in the LMM (p=10*^−^*^8^) but not in any BSLMM run. Using DAPC, 37 SNPs across 26 chromosomes were identified as sex-associated in white suckers (Table S8). A SNP loading statistical threshold of 0.999 was used. This was higher than the threshold used for bluehead suckers to account for the larger number of SNPs and geographic variability. Additionally, there were three F*_ST_* outliers in white suckers based on Rosner’s test, located on chromosomes 1, 24, and 25. Chromosome 1 was identified by all 4 methods (LMM, BSLMM, DAPC, F*_ST_*, Fig.4). Although chromosome 1 was identified by all 4 methods, the SNPs were not the same across the chromosome. Additionally, comparing the sex-associated loci identified by the 4 methods indicates the variability in results for white suckers (Fig. 4).

**Figure 4:**
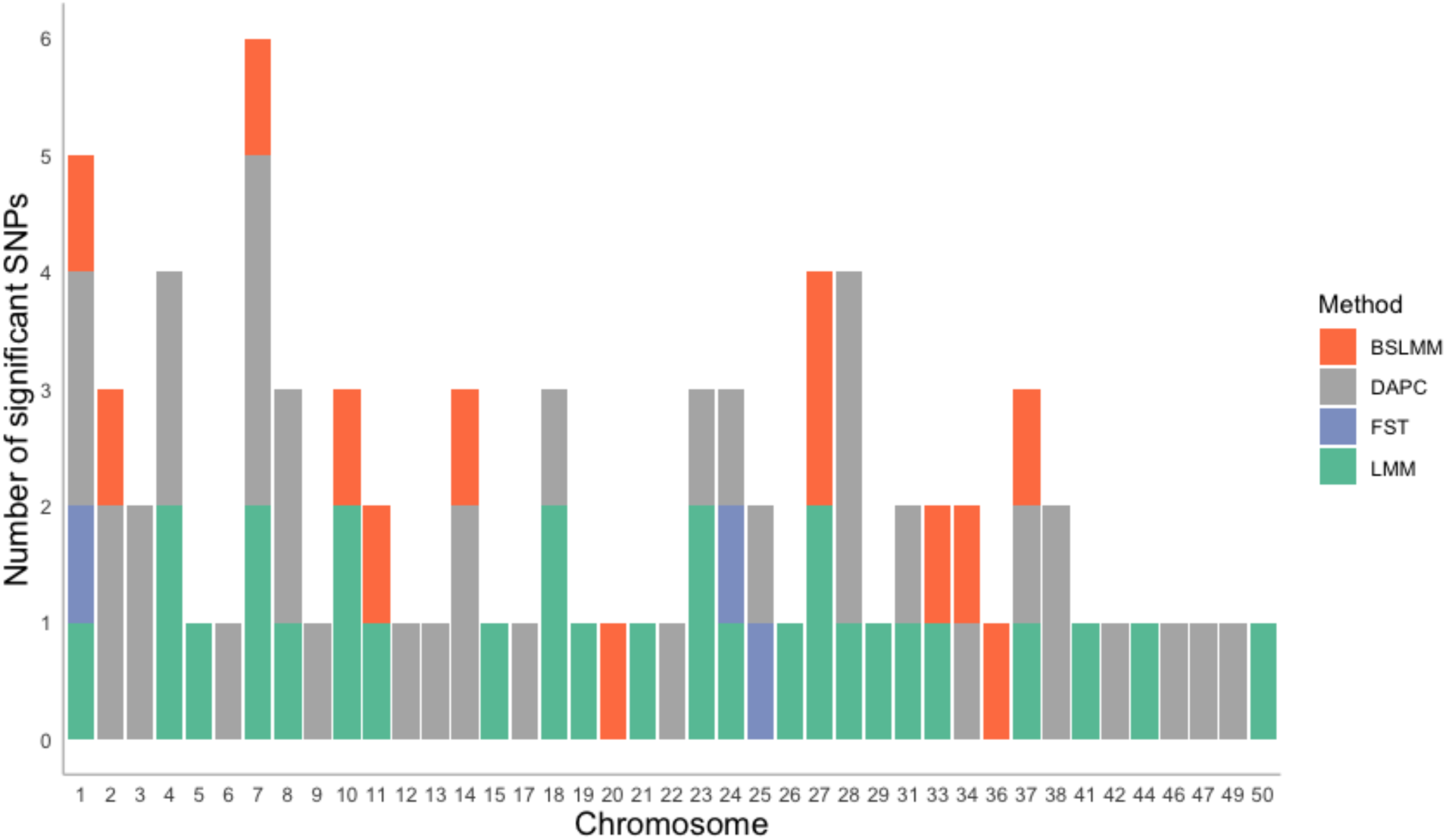
Barplot depicting the overlap of significant SNPs for white suckers (n=44) using GEMMA’s LMM, BSLMM, DAPC, and FST outliers. Only the chromosomes that had significant SNPs are shown on the plot. Only one chromosome has been identified as sex-associated by all 4 methods; however, the sex-associated SNPs are not all the same across methods.

Randomly subsetting the bluehead and entire datasets to explore the effects of differing sample size substantially impacted the LMM and BSLMM results suggested that sample size may have influenced the analyses of white and bluehead×white suckers. For the downsampled bluehead sucker data set, there were no significant loci in any of the three repetitions of the LMM (Fig. S8). With the BSLMM, this pattern was seen in all repetitions (Fig. S9). Interestingly, chromosome 4 was not significant in any run with the reduced bluehead sucker data set. For the reduced data set of all 260 individuals, the same pattern held for the LMM and BSLMM, with none of the repetitions including significant loci (Fig. S10, Fig. S11).

In contrast to bluehead suckers, sex-specific loci (those that are in exclusively in one sex) were detected in white suckers and bluehead×white hybrids using `Ms.SSLI`. There were a total of 27 sex-specific loci found in white suckers (Table S9). These loci were spread across 6 different chromosomes rather than one single chromosome, thus; it is unlikely that a differentiated sex chromosome is present. However, there were two chromosomes with more than 3 sex-specific loci. These were chromosomes 13 and 17. One sex-specific locus was found in bluehead×white hybrids on chromosome 40. While sex specific loci were detected using `Ms.SSLI`, no sex-specific loci were detected using RADSex (Feron *et al*. 2020) (Fig. S5). Bluehead×white hybrids did not have any significant loci in the LMM or any of the BSLMM runs (Fig. S6C, D).

## 5 Discussion

Despite the ubiquity of genetic sex determination and gonochorism in nature, the identity and genetic architecture of sex determination is poorly characterized in many taxa, particularly in groups like fish where there is high evolutionary turnover of sex determination systems (Bachtrog *et al*. 2014, Pennell *et al*. 2018). Given the out-sized role that sex chromosomes and master sex-determining loci potentially play in evolutionary diversification (Heule *et al*. 2014), understanding mechanisms of sex determination is especially desirable in species-rich clades with ongoing interspecific hybridization. This study investigated the genetic basis of sex determination in two species and one hybrid of *Catostomus* fishes using a genome-wide association study and a novel analysis to detect sex-specific loci, `Ms.SSLI`. We present evidence of a genetic basis of sex determination in *Catostomus* fishes, but the exact genetic architecture remains uncertain, and it is unclear how conserved sex determination is across *Catostomus* species. We have found compelling evidence of a sex-determining region on chromosome 4 that explains variation in sex in bluehead suckers. We have also found a significant locus in all species grouped together, but this is most likely attributed to bluehead suckers as they make up more than half of the sample size (Korte & Farlow 2013). Thus, it is unlikely that all species of *Catostomus* suckers have a shared basis of sex determination on chromosome 4; however, it is still possible that they share other sex determining loci. Additionally, different loci are implicated in white suckers, although it is unclear how robust this finding would be to analyses with larger sample sizes. Our work complements the body of research dedicated to understanding how reproductive isolation and hybridization outcomes vary in *Catostomus* fishes. Our study suggests that sex determination may vary across closely related taxa. If there is indeed variation in sex determination mechanisms, the degree to which the genetic architecture of sex determination is shared across species could potentially inhibit or facilitate hybridization in *Catostomus* fishes.

### Evidence of a sex-associated region in bluehead suckers

We found a significant region linked to sex on chromosome 4 in bluehead suckers that is suggestive of an autosomal master sex-determining region (Fig. 3). This chromosome was significant in both the LMM and BSLMM, with large effect sizes. For example, an individual that is homozygous for the significant SNP on chromosome 4 in bluehead suckers would be 67% more likely to be female than male. Previous work on Catostomidae karyotypes, which includes bluehead suckers, indicates the absence of well-differentiated sex chromosomes (Uyeno & Smith 1972). Therefore, it is likely that sex determination is controlled by autosomal sex determination, where a master sex-determining region or regions are present, or by homomorphic sex chromosomes. Beyond the previous karyotype data, our results concur that it is unlikely that there are heteromorphic sex chromosomes because a region as large and differentiated as a heteromorphic sex chromosome would have been detected with RADSex or `Ms.SSLI` (Feron *et al*. 2020). Teleost fish have a wide variety of sex determination mechanisms (Mank *et al*. 2006, Bachtrog *et al*. 2014); however, it is common to have a single sex-determining region control sex differentiation (Charlesworth & Mank 2010, Kongsstovu *et al*. 2020). Further, autosomal sex determination is thought to be ancestral to well-differentiated sex chromosomes (Traut & Winking 2001, Mank *et al*. 2006). Therefore, it is likely that the two significant loci are master sex-determining loci.

The sex-associated region on chromosome 4 could be a precursor to homomorphic sex chromosomes as we found no evidence of heteromorphic sex chromosomes. The potential sex-determining region on chromosome 4 is small as there were only 2 sex-associated variants out of 418; therefore, a small and young homomorphic sex chromosome system could be present (Ogata *et al*. 2021). Sex chromosomes usually evolve from homomorphic to heteromorphic (Charlesworth *et al*. 2005). If the sex chromosomes or regions of *Catostomus* suckers are young, it would be likely that they would take a homomorphic form. In this case, the sex determination region is either young and not diverged, or there has been a change recently in the location of the sex-determining region (Charlesworth & Mank 2010). However, without recombination information and a larger sample size, it is difficult to pinpoint the age and development of possible sex chromosomes.

### Assessing the likelihood of a polygenic architecture in white suckers

We found multiple chromosomes with significantly sex-associated loci in white suckers, potentially indicating a polygenic basis of sex determination (Fig. 2C). However, the smaller sample size of white sucker individuals (n=44) undermines our confidence in a polygenic basis of sex determination to some extent, so caution is warranted in the interpretation of these analyses. BSLMMs are better suited than LMMs for small datasets (Lee & Song 2004, Baldwin & Fellingham 2013), as BSLMMs are Bayesian and thus estimate parameters from data, rather than defining them a priori. These informative parameters and distributions are a better fit to data than those defined a priori (Zhou *et al*. 2013, van de Schoot *et al*. 2015). Thus, the results from the BSLMM model provide more sensitive interpretations of the genetic architecture of sex determination, even with a small sample size. When we down-sampled the entire data set and blueheads to a similar number of individuals to white suckers to explore the effects of sample size, the genetic architecture of sex determination also appeared polygenic, similar to white suckers. This indicates that while the polygenic architecture we detect here may hold with larger sample size, it is also possible that there may be one specific sex-determining region in white suckers (similar to the region on chromosome 4 in bluehead suckers) that cannot be detected due to the sample size. This is an area that merits further work, as larger sample sizes of white suckers and flannelmouth suckers can help clarify whether these three species, which experience extensive contemporary hybridization, have a conserved or variable basis of sex determination.

### Limitations

Certain aspects of our data set may have limited inference in this study. In particular, despite sampling thousands of loci genome-wide, genotyping-by-sequencing typically only samples about 1-2% of the genome which may bias our ability to detect a sex-linked region if the causal loci were not directly sampled, as is likely with this sort of reduced-representation data set (Lowry *et al*. 2017). Some or all of the SNPs that were sex-associated in our data set may not be within the causal locus, but instead might be linked to causal loci. One additional caveat is that it is possible that a female heterogametic (ZW) system with heteromorphic sex chromosomes would be missed because the fish sampled for the reference genome was male, so sequenced genome fragments from a W-like chromosome would not align to our reference genome. This is not just unique to our study, but a consideration that other researchers should take into account. However, regardless of these limitations, our data support the conclusion that the basis of sex determination is genetic, and that loci on chromosome 4 are likely sex-associated in bluehead suckers.

Although evidence supports a genetic basis of sex determination in suckers, we cannot discount the potential for a contributing environmental factor. This is especially the case in white suckers, where there are multiple sex-associated loci. In fish without well-formed sex chromosomes it is often the case that sexual differentiation occurs later in development, which can allow for more environmental influence (Mank *et al*. 2006). It has even been shown that the reproductive responses of white suckers can change in response to lake acidity (Trippel & Harvey 1987). In particular, sexual maturity was found to be later in white suckers in acid lakes in comparison to lakes with more neutral pH, offering a hint that environmental factors may play a role. Only a small fraction of Cypriniformes have been shown to have environmental sex determination (Ashman *et al*. 2014), but closer study of sex determination in other fish lineages has revealed examples of a polygenic basis of sex determination that may interact with environmental variables like temperature (as in European sea bass; Palaiokostas *et al*. 2015). We therefore cannot rule out environmental contributions to sex determination in *Catostomus* fishes, and further study should investigate potential environmental factors.

### Implications of variation in sex-determining mechanisms in *Catostomus* sucker species

The sex-associated loci identified on chromosome 4 in bluehead suckers are potentially part of (or adjacent to) a master sex determining region. However, these loci had little predictive power for sex in white suckers and bluehead×white hybrids, suggesting that bluehead and white suckers may have a different genetic basis of sex determination. Despite potential limitations associated with the sample size of white suckers, it is unlikely that chromosome 4 is sex-associated in white suckers as no loci on chromosome 4 were present in any of the three BSLMM runs with a high PIP or effect size. Other SNPs from chromosome 4 were identified by DAPC and LMM, but neither contained SNPs that were significant in bluehead suckers. Therefore, based on the data presented here, it is unlikely that there is a shared basis of sex determination among *Catostomus* sucker species. Differing sex determination systems could contribute to reproductive isolation between species. Bluehead and white suckers do hybridize in many geographic locations, suggesting that different forms of sex determination are insufficient to fully prevent admixture and are compatible on some level, but at the same time few hybrids beyond the first generation have been observed in this cross (Mandeville *et al*. 2017), which might be indicative of incompatibilities or low fitness of the first generation hybrids. A previous study also suggests asymmetry in ancestry of parents, where white sucker females and bluehead sucker males produce the vast majority of bluehead×white hybrid offspring (Song 2013). It is possible that other factors contribute to reproductive isolation as well, including pre- or post-zygotic factors and genetic or environmental interactions.

Variation in the underlying mechanisms of sex determination may influence reproductive isolation between species (Ser *et al*. 2010, Dufresnes *et al*. 2020, Kottler *et al*. 2020), between populations of the same species (Faggion *et al*. 2019), and even in cytotypes of the same species (Beaudry *et al*. 2020), as sex chromosomes can play a role in the formation and maintenance of post-zygotic isolation. *Catostomus* suckers have variable levels of reproductive isolation both between species and geographically (Mandeville *et al*. 2015, 2017). Bluehead×white hybrids are most often first generation hybrids, with occasional but limited backcrossing when compared to flannelmouth×white hybrids that have extensive spatial variability in hybridization, with more common backcrossing and later generation hybridization (Mandeville *et al*. 2017). The observed differences in the loci contributing to sex between white and bluehead suckers may explain the variation in hybridization between white and bluehead suckers. In particular, the absence of chromosome 4 as a sex-associated region in white suckers may play a role in the limited backcrossing seen in their hybrids.

### Linking genomics to sex assignments

An initial goal of this work was to enable identification of *Catostomus* sucker sex from genetic data alone, to provide context for work on *Catostomus* hybridization. Sex assignments are an important aspect of field work and conservation; however, this study demonstrates that caution should be used when assigning sex using genomic data. When we attempted to predict the phenotype of known individuals based on genetic data, we predicted the correct sex 95% of the time. In contrast, when predicting the sex of individuals of unknown sex, “probable females”, most of the predictions centered around 0.5, indicating sex could not be conclusively determined. This suggests that the genomic regions that are identified using a GWAS or similar method are not enough to assign sex to individuals. However, the number of SNPs controlling sex and specific genetic architecture of sex determination most likely have a large influence on this finding (Trenkel *et al*. 2020), but nonetheless these results have implications for conservation and field work. Studies may use the results from investigations of sex determination for the assignment of sex for unknown individuals (Galindo *et al*. 2011, Trenkel *et al*. 2020, Mussmann *et al*. 2021); we recommend that the predictive power of sex associated markers be explicitly evaluated before those markers are used to determine sex of un-sexed individuals, especially when this information may be used for conservation decisions.

### Conclusion and future directions

In this study, we identified a genetic basis of sex determination in *Catostomus* fishes, and results indicate that the genetic basis of sex determination might vary across species. We located a potential master sex-determining locus on chromosome 4 in bluehead suckers which may play a role in the level of reproductive isolation with white suckers. Although our current results suggest polygenic sex determination in white suckers and bluehead×white hybrids, the identity of the causal loci and regions of the genome remains unclear, and it is possible that sample size limited our statistical power and therefore the scope of our inferences. While the full role of sex-determination systems in reproductive isolation remains unclear, we can speculate that sex determination might exert some influence on the level of reproductive isolation seen between different *Catostomus* sucker species. Additional work with these species, including bolstering sample sizes, including additional species (especially flannelmouth), and sequencing a larger fraction of the genome, will reveal more fully the extent to which sex determination loci might affect current hybridization dynamics and what role sex determination has played in evolutionary diversification of the Catostomidae.

## Data accessibility

Upon acceptance of this manuscript, all scripts and data will be uploaded to Data Dryad, NCBI SRA, and Github. Data can also be made available to reviewers upon request.

## Author contributions

CBP and EGM planned research, designed analysis strategy, generated genomic data, and implemented bioinformatics workflows. SEM contributed to analyses. All authors wrote and revised the manuscript.

## Acknowledgements

We would like to thank collaborators for assistance in acquiring fin tissues, specifically the McCann lab at the University of Guelph, Colorado Parks and Wildlife, and the Wyoming Game and Fish Department. This work was funded by an NSERC Discovery Grant, the Canada First Research Excellence Fund Food From Thought program at the University of Guelph, and startup funding from the University of Guelph to E.G. Mandeville. S.E. McFarlane was supported by the *modelscape* project with funding from NSF award number 2019528. Computing was completed using a RRG Allocation on Compute Canada’s Graham cluster to E.G. Mandeville. We would also like to thank Amanda Meuser, Jill Campbell, Ben Schultz, and Amy Pitura for their comments and discussion on earlier drafts of this manuscript.

