## Supplement for "Identification of sex-determining loci in hybridizing *Catostomus* fish species"

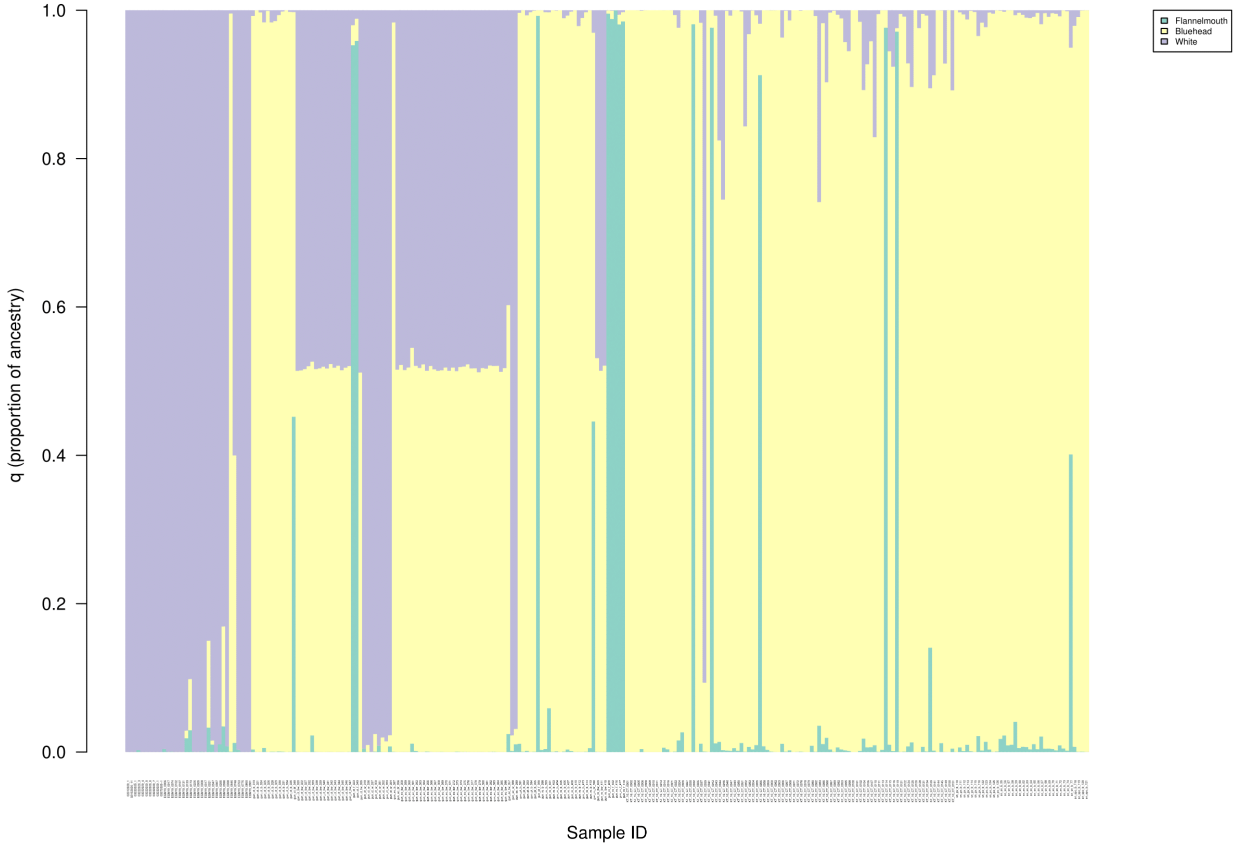


Figure S1: Estimates of q (proportion of ancestry) for all individuals used in the study after filtering (n=260) with k=3. Each individual is listed on the x axis, with its proportion of ancestry on the y axis. Each of the 3 species is coloured and corresponds to the legend.

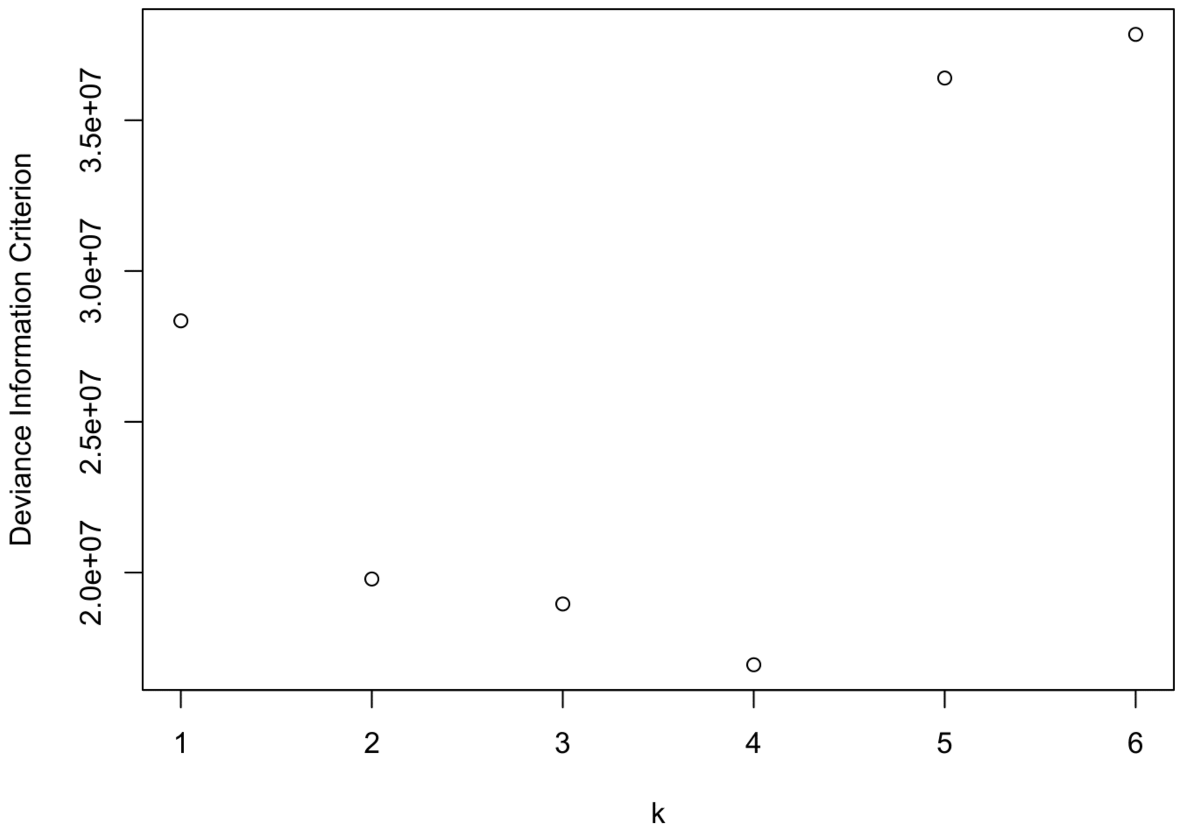


Figure S2: DIC (Deviance information criterion) values from entropy for each value of k.

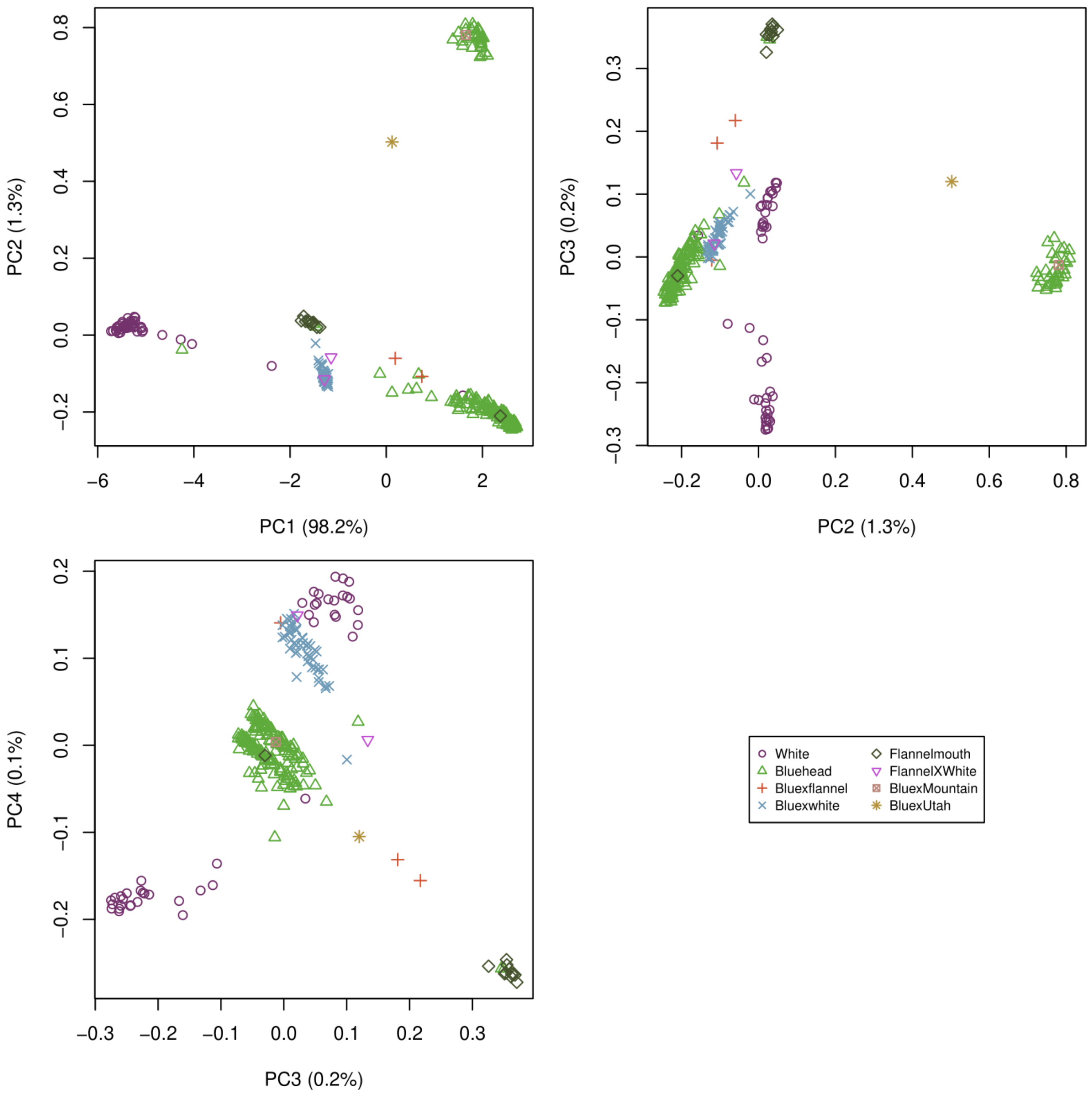


Figure S3: Principal component analysis for bluehead, flannelmouth, and white suckers as well as their hybrids. Each point represents an individual fish based on phenotypic identification. The principal components separate by species. PC1 separates the parental species from each other. PC2 separates the bluehead suckers by geography.

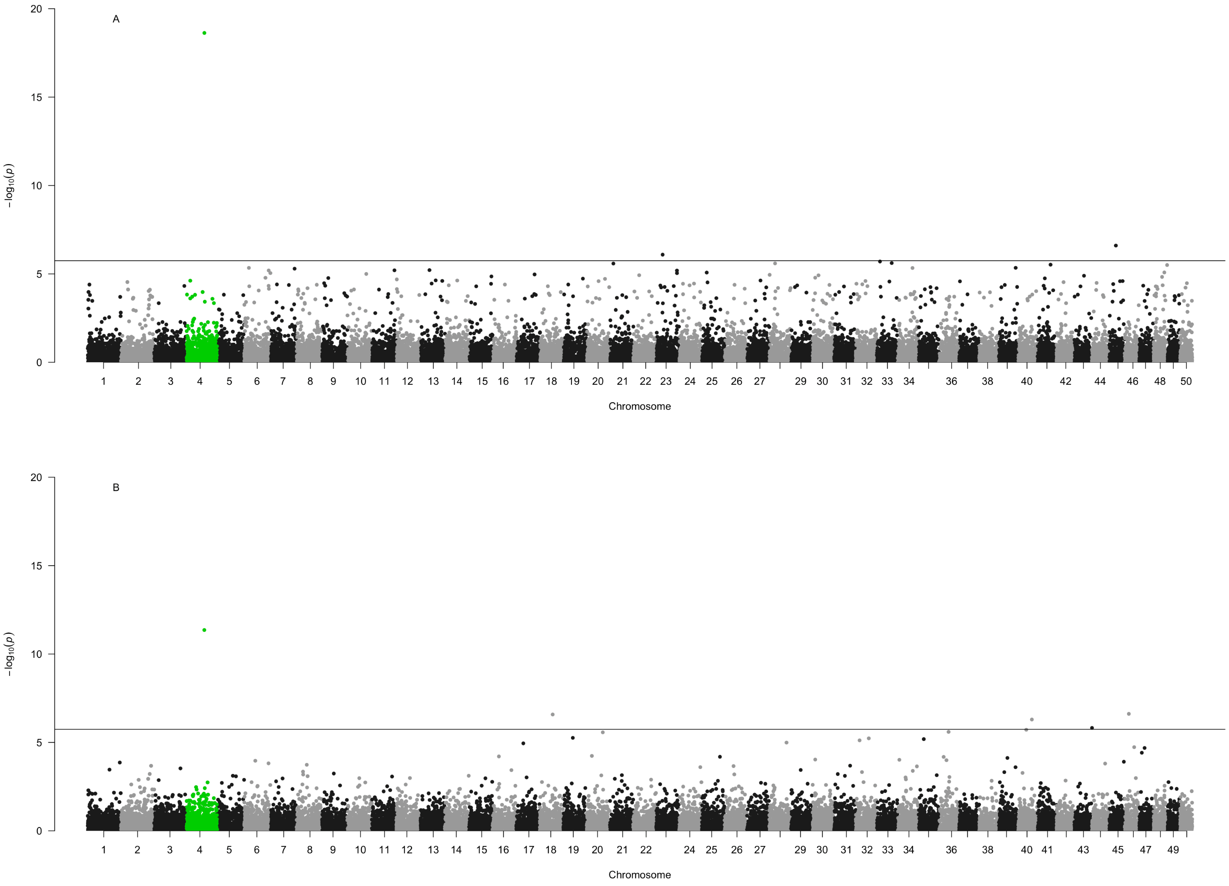


Figure S4: Manhattan plots showing the difference between the datasets with and without 'probable females'. a). Data set (n=260) with 'probable females' included as females. b). Data set (n=211) with 'probable females' excluded.

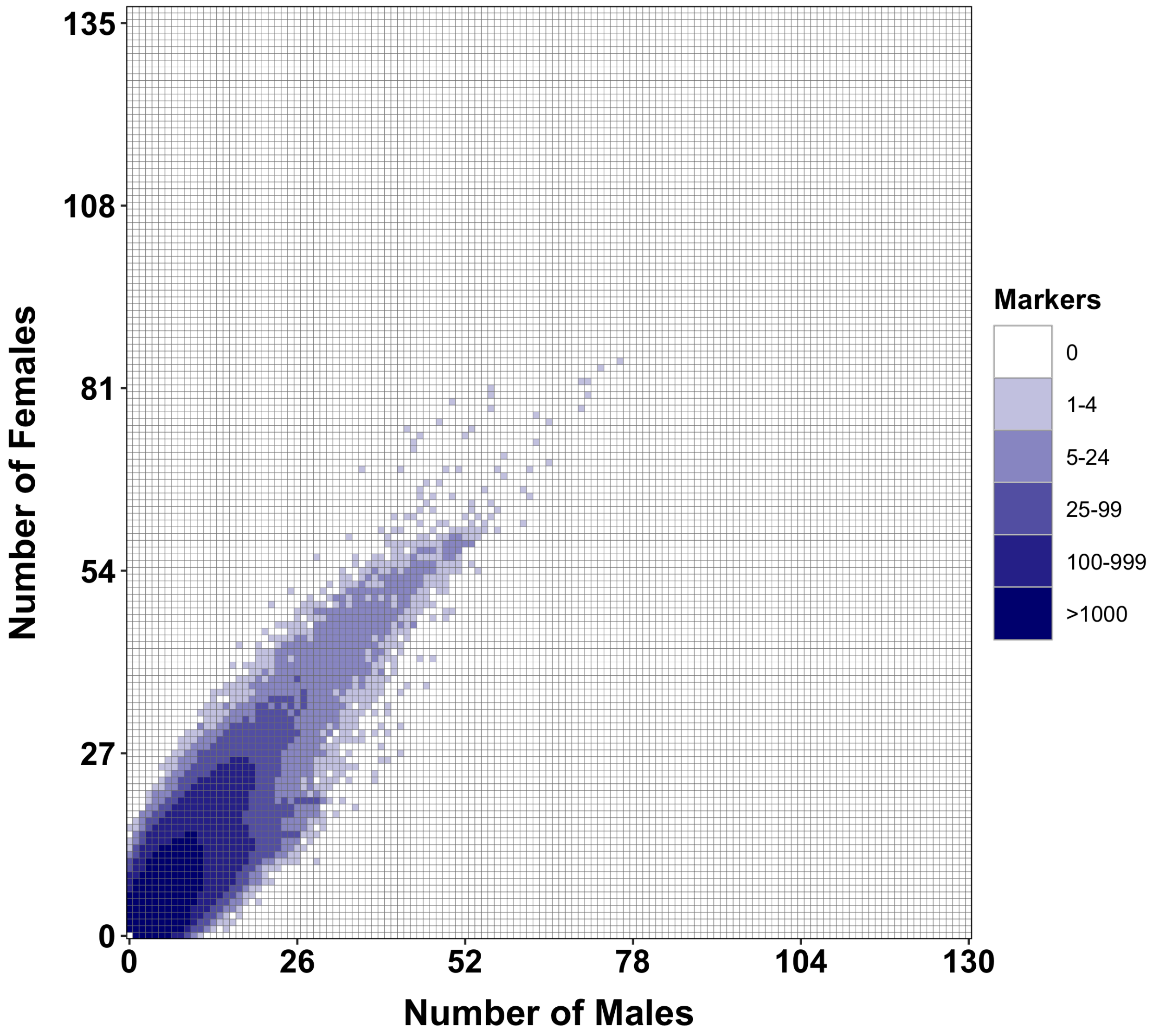


Figure S5: Tile plot of potential sex-specific markers from RadSex from the full data set (n=260). The colour of the square indicates the number of shared markers in males and females. If markers were significantly found in one sex then the box would be surrounded by red; thus, there were no sex-specific markers found.

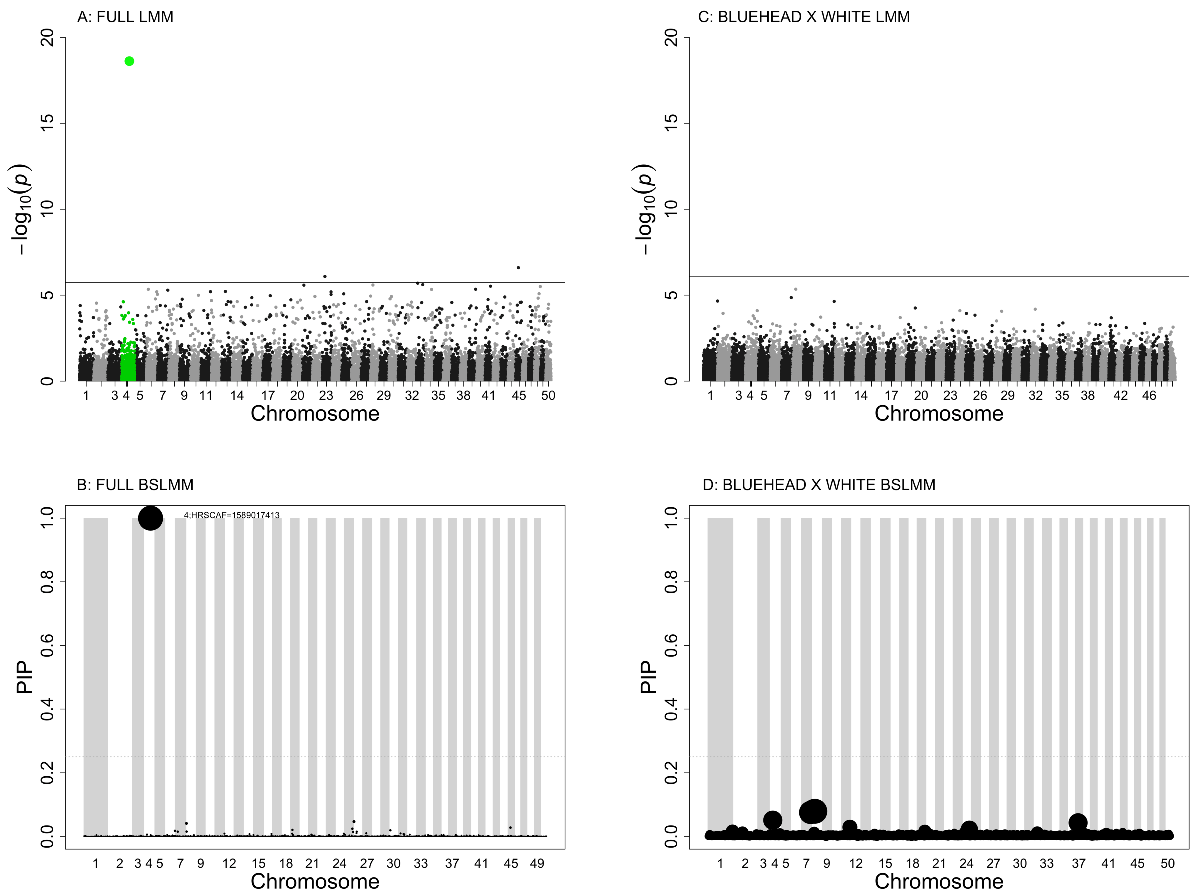


Figure S6: Sex-associated loci in the entire *Catostomus* and bluehead×white hybrids across the 50 largest scaffolds relative to the flannelmouth reference genome. a) Manhattan plot from GEMMA's LMM between SNPs and sex for the entire dataset (n=260). The black line indicates the Bonferonni corrected significance threshold of $p=7.20^{-7}$. b) PIP of SNPs from GEMMA's BSLMM for the entire sucker data set (n=260). The dotted line indicates the threshold of PIP value of 0.25, and the grey and white blocks visually show the chromosomes. The sparse effect is indicated by the size of the point. c) Manhattan plot from GEMMA's LMM between SNPs and sex for bluehead×white hybrids (n=46), with a significance threshold of $p=8.43^{-7}$. d) PIP of SNPs across chromosomes from the BSLMM for bluehead×white hybrids. As with b), The dotted line indicates the threshold of PIP value of 0.25 and the sparse effect is indicated by the size of the point.

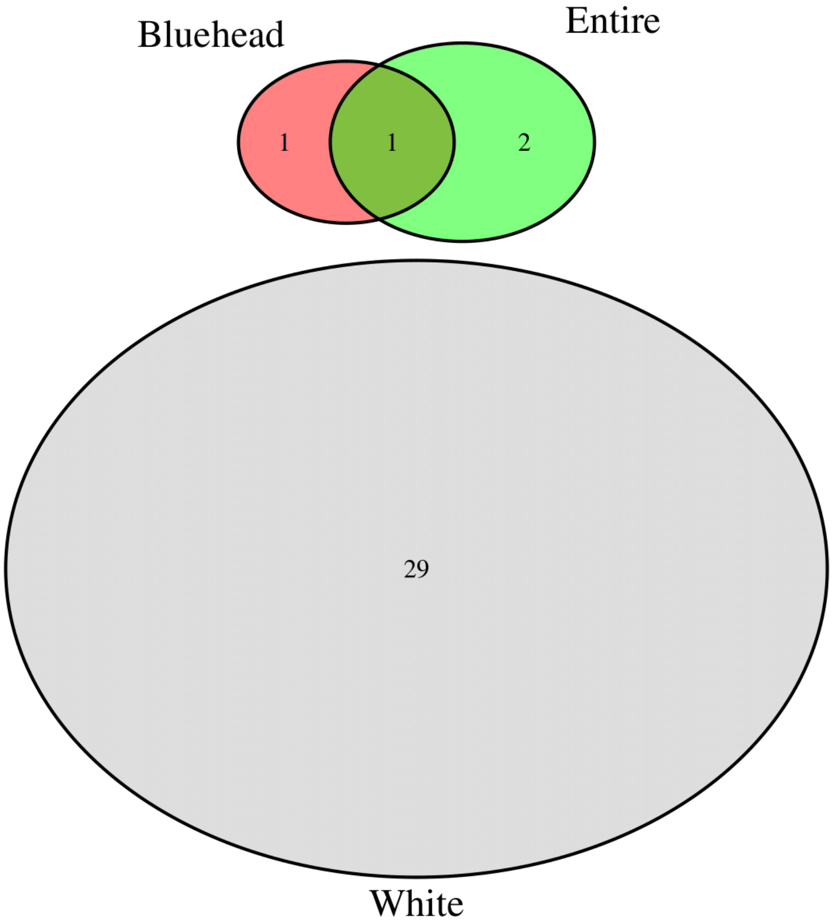


Figure S7: Venn diagram showing the overlap of significant SNPs from LMM between the entire dataset (n=260), bluehead (n=150), and white (n=44) suckers.

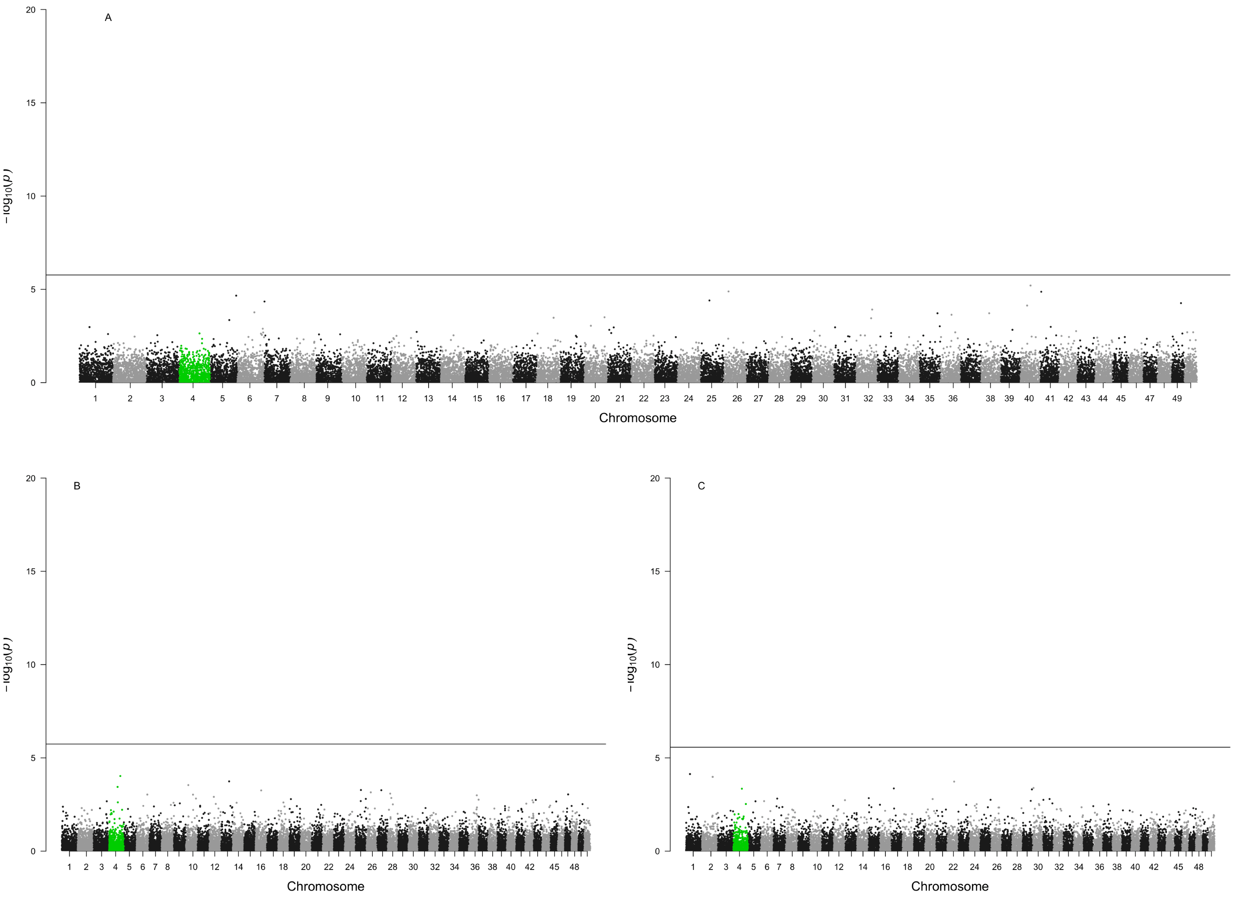


Figure S8: Manhattan plots showing the results for the three LMM down-sampled repetitions for the bluehead (n=150) data set.

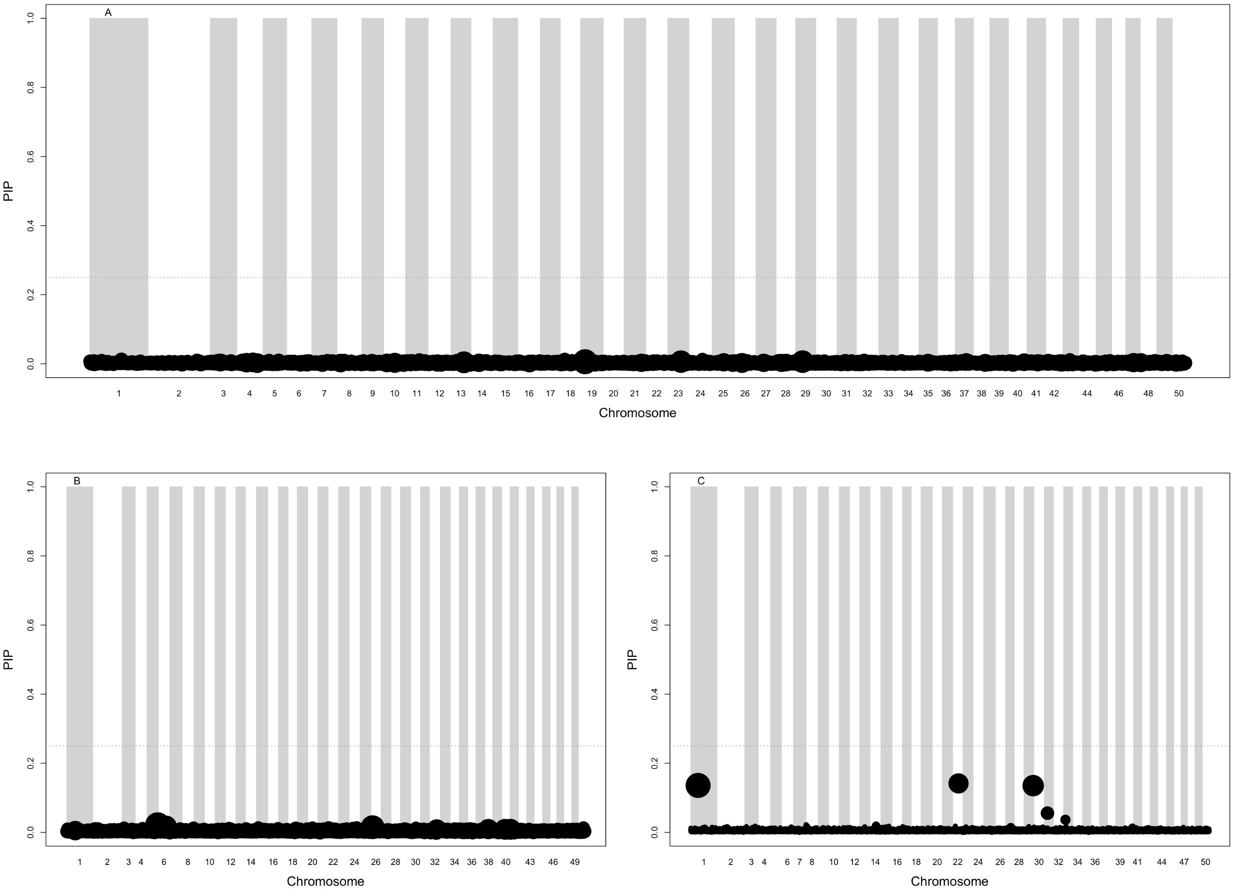


Figure S9: Manhattan plots showing the results for the three BSLMM down-sampled repetitions for the bluehead (n=150) data set.

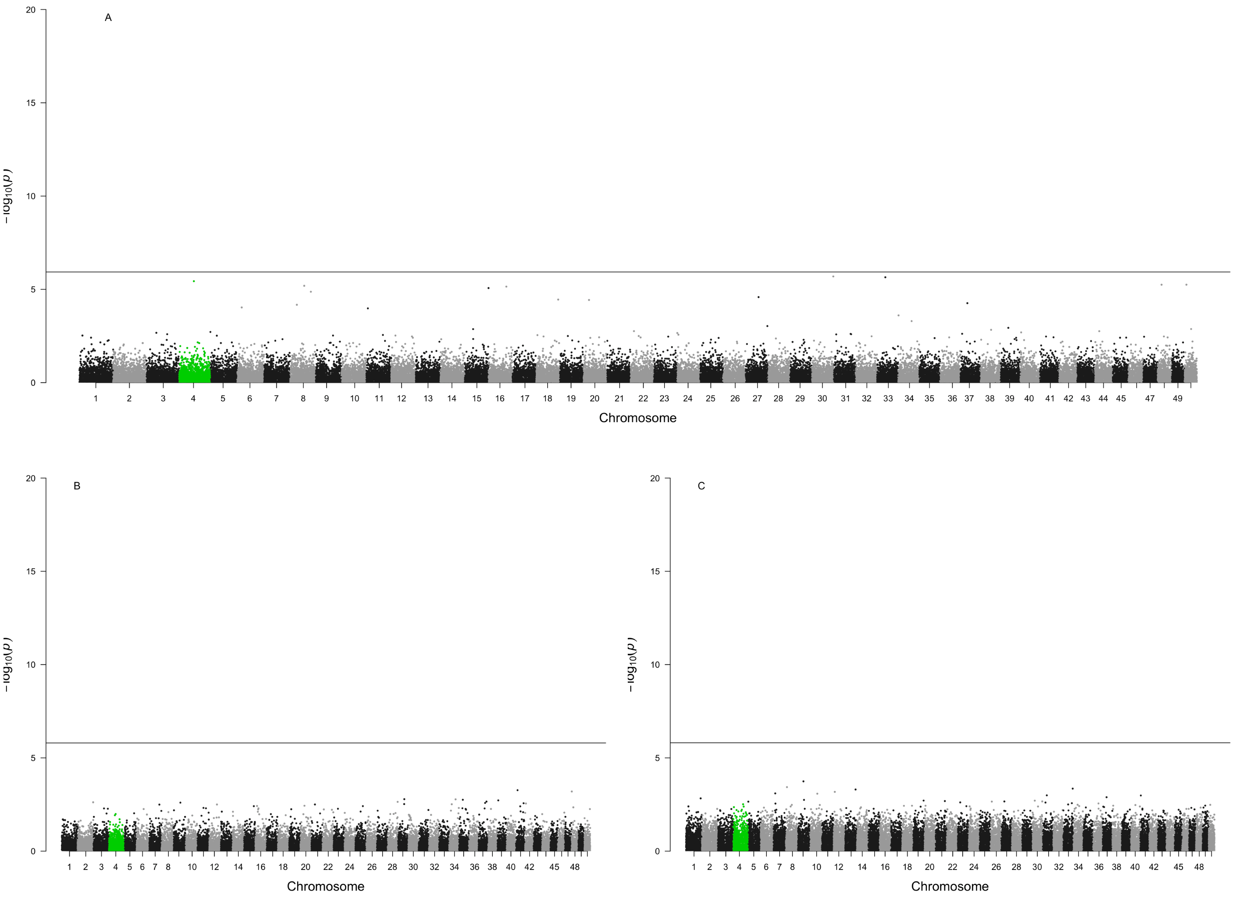


Figure S10: Manhattan plots showing the results for the three LMM down-sampled repetitions for the full (n=260) data set.

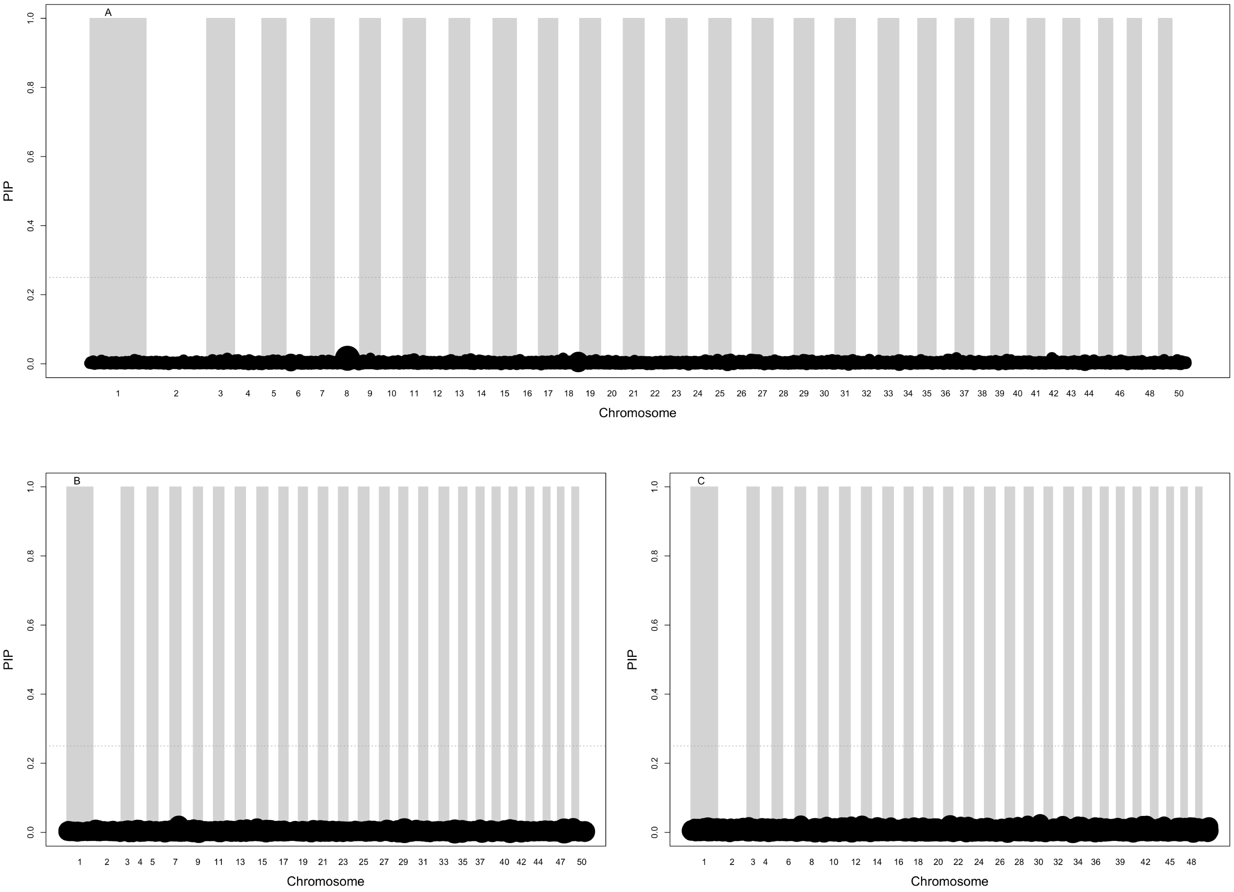


Figure S11: Manhattan plots showing the results for the three BSLMM down-sampled repetitions for the full (n=260) data set.

Table S1: Number of individuals per species or hybrid and location in this study. There are a total of 269 individuals sampled from Colorado, Ontario, and Wyoming. The total number from each sampling location can be seen across the columns and the total number of individuals from each species or hybrid can be seen across the rows.

| **Species** | **Colorado** | **Ontario** | **Wyoming** | **Total** |
|---|---|---|---|---|
| White sucker | 20 | 26 | 0 | 46 |
| Flannelmouth sucker | 12 | 0 | 0 | 12 |
| Bluehead sucker | 121 | 0 | 34 | 155 |
| Bluehead×flannelmouth hybrid | 3 | 0 | 0 | 3 |
| Bluehead×white hybrid | 47 | 0 | 1 | 48 |
| Flannelmouth×white hybrid | 2 | 0 | 0 | 2 |
| Mountain x white hybrid | 0 | 0 | 1 | 1 |
| Bluehead x mountain hybrid | 0 | 0 | 1 | 1 |
| Bluehead x Utah hybrid | 0 | 0 | 1 | 1 |
| Total | 205 | 26 | 38 | **269** |

Table S2: Number of individuals per species or hybrid and their sex. Males, females and probable females (those that could not be conclusively sexed as female) are included.

| **Species** | **Males** | **Females** | **Probable Females** | **Total** |
|---|---|---|---|---|
| White sucker | 33 | 9 | 4 | 46 |
| Flannelmouth sucker | 7 | 3 | 2 | 12 |
| Bluehead sucker | 73 | 37 | 45 | 155 |
| Bluehead×flannelmouth hybrid | 1 | 2 | 0 | 3 |
| Bluehead×white hybrid | 15 | 33 | 0 | 48 |
| Flannelmouth×white hybrid | 1 | 1 | 0 | 2 |
| Mountain x white hybrid | 1 | 0 | 0 | 1 |
| Bluehead x mountain hybrid | 0 | 1 | 0 | 1 |
| Bluehead x Utah hybrid | 0 | 1 | 0 | 1 |
| Total | 131 | 87 | 51 | **269** |

Table S3: SNPs that have a PIP value of 0.25 or higher indicating that they were included in the sparse distribution at least 25 percent of the time for bluehead suckers (n=150), white suckers (n=44) and the full data set (n=260).

| | | | BSLMM RUN A | | BSLMM RUN B | | BSLMM RUN C | |
|---|---|---|---|---|---|---|---|---|
| **Species** | **Chromosome** | **Base pair position** | **PIP** | **Effect size** | **PIP** | **Effect size** | **PIP** | **Effect size** |
| Bluehead | 4;HRSCAF=15890 | 27210425 | 1 | 0.6 | 1 | 0.61 | 1 | 0.61 |
| Bluehead | 19;HRSCAF=14438 | 14763515 | 0.35 | 0.15 | 0.38 | 0.13 | 0.32 | 0.14 |
| Bluehead | 4;HRSCAF=15890 | 27220056 | 0.42 | 0.11 | 0.38 | 0.1 | 0.38 | 0.1 |
| Full | HRSCAF=15890 | 27210425 | 1 | 0.61 | 1 | 0.61 | 1 | 0.61 |
| White | 2;HRSCAF=15409 | 42071626 | 1 | 0.96 | - | - | - | - |
| White | 20;HRSCAF=14974 | 31322275 | 1 | 0.93 | - | - | - | - |
| White | 34;HRSCAF=14797 | 28590989 | 1 | 0.92 | - | - | - | - |
| White | 27;HRSCAF=15177 | 12966734 | - | - | 1 | 0.98 | - | - |
| White | 27;HRSCAF=15177 | 4750078 | - | - | - | - | 1 | 0.49 |
| White | 11;HRSCAF=13688 | 19396195 | - | - | 1 | 0.96 | - | - |
| White | 7;HRSCAF=15635 | 11503782 | - | - | 1 | 0.95 | - | - |
| White | 10;HRSCAF=15274 | 29119186 | - | - | - | - | 1 | 0.52 |
| White | 14;HRSCAF=16429 | 9089851 | - | - | - | - | 1 | 0.37 |
| White | 37;HRSCAF=16427 | 9461853 | - | - | - | - | 1 | 0.31 |
| White | 1;HRSCAF=16083 | 6548878 | - | - | - | - | 1 | 0.22 |
| White | 33;HRSCAF=6958 | 1784666 | - | - | - | - | 1 | 0.17 |

Table S4: Significant SNPs in the linear mixed model for bluehead suckers (*C. discobolus*) (n=150). Significance was determined using the Bonferroni correction. We report the chromosome and SNP ID for each significant SNP as well as the p-value and beta estimate as a proxy for the effect size.

| **Chromosome** | **Base pair position** | **P-value** | **Beta estimate** |
|---|---|---|---|
| 4;HRSCAF=15890 | 27210425 | $10^{-15}$ | 0.67 |
| 4;HRSCAF=15890 | 27320056 | $10^{-10}$ | 0.43 |

Table S5: Significant SNPs in the linear mixed model for the full data set (n=260). Significance was determined using the Bonferroni correction. We report the chromosome, SNP ID, and base pair position for each significant SNP as well as the p-value and beta estimate as a proxy for the effect size.

| **Chromosome** | **Base pair position** | **P-value** | **Beta estimate** |
|---|---|---|---|
| 4;HRSCAF=15890 | 27210425 | $10^{-19}$ | 0.63 |
| 23; HRSCAF=16445 | 9380921 | $10^{-7}$ | 0.33 |
| 45;HRSCAF=5992 | 10211305 | $10^{-7}$ | 0.27 |

Table S6: Significant SNPs in the linear mixed model for white suckers (*C. commersonii* (n=44). Significance was determined using the Bonferroni correction. We report the chromosome and base pair position for each significant SNP as well as the p-value and beta estimate as a proxy for the effect size.

| **Chromosome** | **Base pair position** | **P-value** | **Beta estimate** |
|---|---|---|---|
| 1;HRSCAF=16083 | 16272483 | $10^{-7}$ | 0.89 |
| 4;HRSCAF=15890 | 19023513 | $10^{-8}$ | 0.63 |
| 4;HRSCAF=15890 | 38654147 | $10^{-7}$ | 0.74 |
| 5;HRSCAF=13969 | 12878581 | $10^{-7}$ | 0.47 |
| 7;HRSCAF=15635 | 23872043 | $10^{-7}$ | 0.53 |
| 7;HRSCAF=15635 | 31194957 | $10^{-7}$ | 0.57 |
| 8;HRSCAF=13978 | 2668407 | $10^{-7}$ | 0.51 |
| 10;HRSCAF=15274 | 6599259 | $10^{-8}$ | 0.62 |
| 10;HRSCAF=15274 | 35578887 | $10^{-7}$ | 0.86 |
| 11;HRSCAF=13688 | 37763697 | $10^{-7}$ | 0.52 |
| 15;HRSCAF=4813 | 4368412 | $10^{-7}$ | 0.47 |
| 18;HRSCAF=16431 | 13043088 | $10^{-9}$ | 0.52 |
| 18;HRSCAF=16431 | 21388714 | $10^{-7}$ | 0.56 |
| 19;HRSCAF=14438 | 6235904 | $10^{-7}$ | 0.51 |
| 21;HRSCAF=4986 | 13078704 | $10^{-12}$ | 0.89 |
| 23;HRSCAF=16445 | 9116100 | $10^{-7}$ | 0.68 |
| 23;HRSCAF=16445 | 16749748 | $10^{-7}$ | 0.89 |
| 24;HRSCAF=13971 | 32967439 | $10^{-7}$ | 0.37 |
| 26;HRSCAF=5332 | 18691305 | $10^{-7}$ | 0.65 |
| 27;HRSCAF=15177 | 4750078 | $10^{-10}$ | 0.57 |
| 27;HRSCAF=15177 | 6062398 | $10^{-9}$ | 0.52 |
| 28;HRSCAF=15710 | 21556660 | $10^{-7}$ | 0.57 |
| 29;HRSCAF=16447 | 8272095 | $10^{-8}$ | 0.80 |
| 31;HRSCAF=111 | 2930809 | $10^{-8}$ | 0.45 |
| 33;HRSCAF=6958 | 24485888 | $10^{-7}$ | 0.68 |
| 37;HRSCAF=16477 | 12195703 | $10^{-7}$ | 0.39 |
| 41;HRSCAF=16049 | 5188100 | $10^{-9}$ | 0.78 |
| 44;HRSCAF=16428 | 22468077 | $10^{-8}$ | 0.73 |
| 50;HRSCAF=16438 | 11768245 | $10^{-11}$ | 0.90 |

Table S7: Sex-associated SNPs from DAPC for bluehead suckers (*C. discobolus*) (n=150). Significance was determined using a SNP loading threshold of 0.995. We report the chromosome, base pair position, and SNP loading for each significant SNP .

| **Chromosome** | **Base pair position** | **SNP loading** |
|---|---|---|
| 1;HRSCAF=16083 | 28983120 | 0.0007733727 |
| 2;HRSCAF=15409 | 10470262 | 0.0008061907 |
| 3;HRSCAF=14022 | 11431534 | 0.0018481672 |
| 4;HRSCAF=15890 | 27320056 | 0.0007421422 |
| 4;HRSCAF=15890 | 27473930 | 0.0007579843 |
| 4;HRSCAF=15890 | 32243401 | 0.0010773495 |
| 5;HRSCAF=13969 | 5920546 | 0.0011861079 |
| 7;HRSCAF=15635 | 36279681 | 0.0009936535 |
| 8;HRSCAF=13978 | 16715685 | 0.0007660276 |
| 10;HRSCAF=15274 | 6821183 | 0.0008947130 |
| 10;HRSCAF=15274 | 37795890 | 0.0014277243 |
| 12;HRSCAF=16439 | 330705 | 0.0010847125 |
| 12;HRSCAF=16439 | 3664944 | 0.0007239448 |
| 12;HRSCAF=16439 | 7495340 | 0.0011029514 |
| 13;HRSCAF=16443 | 2260089 | 0.0007405890 |
| 13;HRSCAF=16443 | 14166300 | 0.0008021539 |
| 14;HRSCAF=16429 | 13179236 | 0.0008252172 |
| 14;HRSCAF=16429 | 18313622 | 0.0008043284 |
| 14;HRSCAF=16429 | 36820402 | 0.0007258998 |
| 15;HRSCAF=4813 | 11988274 | 0.0008925184 |
| 16;HRSCAF=15964 | 10182129 | 0.0011942481 |
| 19;HRSCAF=14438 | 4810057 | 0.0008026318 |
| 19;HRSCAF=14438 | 14121380 | 0.0012275446 |
| 20;HRSCAF=14974 | 14589559 | 0.0007349202 |
| 21;HRSCAF=4986 | 22905624 | 0.0007419157 |
| 23;HRSCAF=16445 | 14564736 | 0.0010979660 |
| 23;HRSCAF=16445 | 29252861 | 0.0016820307 |
| 24;HRSCAF=13971 | 15369346 | 0.0009157206 |
| 25;HRSCAF=14217 | 21585662 | 0.0008108371 |
| 28;HRSCAF=15710 | 1225690 | 0.0007397734 |
| 30;HRSCAF=16430 | 22882372 | 0.0009090379 |
| 31;HRSCAF=111 | 400891 | 0.0007878184 |
| 32;HRSCAF=5184 | 23491073 | 0.0009865368 |
| 32;HRSCAF=5184 | 28097670 | 0.0007389490 |
| 33;HRSCAF=6958 | 3156455 | 0.0007865875 |
| 34;HRSCAF=14797 | 18594159 | 0.0007224144 |
| 34;HRSCAF=14797 | 31451238 | 0.0011164695 |
| 35;HRSCAF=11055 | 6590630 | 0.0008570310 |
| 35;HRSCAF=11055 | 23557785 | 0.0013512806 |
| 35;HRSCAF=11055 | 23971359 | 0.0007790677 |
| 36;HRSCAF=14733 | 19791531 | 0.0008645088 |
| 36;HRSCAF=14733 | 23602599 | 0.0007767997 |
| 36;HRSCAF=14733 | 23722077 | 0.0008811477 |
| 37;HRSCAF=16427 | 5313090 | 0.0009001484 |
| 39;HRSCAF=16448 | 6939402 | 0.0008256468 |
| [illegible] | [illegible] | [illegible] |

Table S8: Sex-associated SNPs from DAPC for white suckers (*C. commersonii* (n=44). Significance was determined using a SNP loading threshold of 0.999. We report the chromosome, base pair position, and SNP loading for each significant SNP .

| **Chromosome** | **Base pair position** | **SNP loading** |
|---|---|---|
| 1;HRSCAF=16083 | 2487552 | 0.0003436758 |
| 1;HRSCAF=16083 | 3845919 | 0.0003377555 |
| 2;HRSCAF=15409 | 11563274 | 0.0002467396 |
| 2;HRSCAF=15409 | 18061014 | 0.0002650946 |
| 3;HRSCAF=14022 | 5680026 | 0.0002561366 |
| 3;HRSCAF=14022 | 50030809 | 0.0002589539 |
| 4;HRSCAF=15890 | 32013696 | 0.0003415403 |
| 4;HRSCAF=15890 | 36145027 | 0.0003397481 |
| 5;HRSCAF=13969 | 5426253 | 0.0003053995 |
| 7;HRSCAF=15635 | 1344462 | 0.0006674428 |
| 7;HRSCAF=15635 | 36267609 | 0.0002645778 |
| 7;HRSCAF=15635 | 36655138 | 0.0002504582 |
| 8;HRSCAF=13978 | 2668407 | 0.0002861939 |
| 8;HRSCAF=13978 | 25036473 | 0.0002754238 |
| 9;HRSCAF=13652 | 34093557 | 0.0002593831 |
| 12;HRSCAF=16439 | 5749521 | 0.0002859795 |
| 13;HRSCAF=16443 | 11308179 | 0.0002740620 |
| 14;HRSCAF=16429 | 14187527 | 0.0002742000 |
| 14;HRSCAF=16429 | 36820391 | 0.0002569439 |
| 17;HRSCAF=14082 | 30015110 | 0.0002813580 |
| 18;HRSCAF=16431 | 6036544 | 0.0003688196 |
| 22;HRSCAF=16435 | 14545145 | 0.0002640777 |
| 23;HRSCAF=16445 | 29252861 | 0.0002477076 |
| 24;HRSCAF=13971 | 10134326 | 0.0002511970 |
| 25;HRSCAF=14217 | 3269474 | 0.0002641016 |
| 28;HRSCAF=15710 | 6068866 | 0.0003717339 |
| 28;HRSCAF=15710 | 20909956 | 0.0003504012 |
| 28;HRSCAF=15710 | 27704991 | 0.0002621973 |
| 31;HRSCAF=111 | 26700847 | 0.0002542989 |
| 34;HRSCAF=14797 | 22421973 | 0.0003299693 |
| 37;HRSCAF=16427 | 14054087 | 0.0003778015 |
| 38;HRSCAF=14493 | 14432602 | 0.0002584134 |
| 38;HRSCAF=14493 | 17076180 | 0.0002744650 |
| 42;HRSCAF=1736 | 6812807 | 0.0002496439 |
| 46;HRSCAF=10957 | 18802739 | 0.0002487577 |
| 47;HRSCAF=16437 | 17736419 | 0.0002458242 |
| 49;HRSCAF=16207 | 4614123 | 0.0003103004 |

Table S9: Sex-specific loci identified using the `Ms.SSLI` program in white suckers (*C.commersonii*). The number of loci identified for each chromosome are listed. Additionally, the base pair position of each of the loci are listed.

| Chromosome | Number of loci | Base pair positions |
|---|---|---|
| 2 | 1 | 40942990 |
| 7 | 1 | 25347653 |
| 10 | 1 | 25601514 |
| 13 | 3 | 2349871, 10369672, 32458127 |
| 17 | 3 | 5286133, 27040337, 32281798 |
| 19 | 2 | 11715482, 35067288 |
| 20 | 1 | 3751390 |
| 21 | 1 | 7546255 |
| 24 | 1 | 8314264 |
| 29 | 1 | 29135399 |
| 32 | 1 | 2374052 |
| 33 | 1 | 29141350 |
| 36 | 1 | 27196854 |
| 38 | 1 | 2383499 |
| 42 | 1 | 26949376 |
| 44 | 2 | 5342499, 20082396 |
| 45 | 2 | 6140859, 13239845 |
| 46 | 1 | 3268477 |
| 50 | 2 | 15779099, 16355689 |

Table S10: Significant SNPs from the linear mixed model for the ten incorrectly sexed permutations (n=211). The permutations did not include probable females.

| Permutation | Chromosome | Base pair positions | P-value | Beta estimate |
|---|---|---|---|---|
| 1 | NA | NA | NA | NA |
| 2 | NA | NA | NA | NA |
| 3 | NA | NA | NA | NA |
| 4 | NA | NA | NA | NA |
| 5 | NA | NA | NA | NA |
| 6 | 36; HRSCAF=14733 | 15052855 | $10^{-9}$ | 0.41 |
| 7 | NA | NA | NA | NA |
| 8 | NA | NA | NA | NA |
| 9 | NA | NA | NA | NA |
| 10 | 4; HRSCAF=15890 | 27210425 | $10^{-7}$ | 0.42 |

### Probable Females

#### Material and Methods

First, we ran the LMM three times: initially, with the 'probable females' removed, then, coded as females, and finally, coded as a third sex to determine whether the inclusion of probable females influenced the results. We also used the LMM to determine whether incorrectly sexed individuals would result in a similar outcome to the data set including probable females. To do this, we removed the 'probable females' and ran 10 permutations where 20 percent of the individuals were mis-sexed and run through GEMMA's LMM. We then used BSLMM and GEMMA's phenotype prediction functionality to predict the phenotype of the probable females. The phenotype prediction uses the output from running BSLMM. The entire data set with 'probable females' was run with the phenotype of 'probable females' as "NA". To determine the predictability of the loci, we also ran the phenotype prediction function with 20 percent of the sexed individuals without sex information (i.e. "NA"). This data set did not include the probable females. Lastly, the third run of the phenotype prediction just included bluehead and white suckers to investigate whether the bluehead suckers' sex-associated loci could be used to predict the sex of white suckers.

#### Results

In both cases, chromosome 4 contained multiple significant loci with not many other loci reaching the significance threshold. In contrast, when we permuted the data ten times and intentionally incorrectly assigned sex to 20 percent of the individuals, we found evidence of a polygenic SD. For the incorrectly sexed permutations, each result from the LMM was quite different to each other and to the entire data set with and without probable females included. Only two of ten permutations resulted in a significant SNP, one of which was on chromosome 4 (Table S10). This indicates that the probable females are females.

GEMMA's phenotype prediction functionality resulted in a range of phenotype values when attempting to predict the sex of the 'probable female'. In a regular phenotype file, males are coded as zeroes and females are coded as ones. In the predicted phenotype file, the predicted phenotypes ranged from 0 to 1. Many values centered around 0.5, therefore indicating that those individuals could not be predicted to be clearly a male or female. To determine whether GEMMA could predict phenotype for known individuals, we removed the probable females and removed sex information for 20 percent of the sexed individuals. The results were still variable; however, the values could be matched with the correct sex 95 percent of the time. This means that 95 percent of the time, if the value was below 0.5 the field-assigned sex was male and if it was above 0.5 the field-assigned sex was female.
